# Circadian and environmental signal integration in a natural population of *Arabidopsis*

**DOI:** 10.1101/2022.09.10.507414

**Authors:** Haruki Nishio, Dora L. Cano-Ramirez, Tomoaki Muranaka, Luíza Lane de Barros Dantas, Mie N. Honjo, Jiro Sugisaka, Hiroshi Kudoh, Antony N. Dodd

## Abstract

Plants sense and respond to environmental cues during 24 h fluctuations in their environment. This requires the integration of internal cues such as circadian timing with environmental cues such as light and temperature to elicit cellular responses through signal transduction. However, the integration and transduction of circadian and environmental signals within plants growing in natural environments remains poorly understood. To gain insights into the 24 h dynamics of environmental signalling in nature, we performed a field study of signalling from the nucleus to chloroplasts in a natural population of *Arabidopsis halleri.* Using advanced modelling approaches to interpret the data, we identified that the circadian clock and temperature are key regulators of this pathway under natural conditions. We identified potential time-delay steps between pathway components, and diel fluctuations in the response of the pathway to temperature cues that are reminiscent of the process of circadian gating. This approach of combining studies of gene expression in the field with modelling allowed us to identify the dynamic integration and transduction of environmental cues, in plant cells, under naturally fluctuating diel cycles.

## Introduction

Plants have sophisticated environmental sensing and signalling mechanisms that underpin their responses to the fluctuating environment. Under natural conditions, this requires signalling pathways that integrate dynamic, overlapping and complex environmental stimuli [1, 2]. These environmental fluctuations include an oscillating component with a 24 h period, which arises from the cycle of day and night. Circadian clocks, which are endogenous biological oscillators, provide a cellular estimate of the time of day that coordinates the diel responses of plants with environmental fluctuations by aligning transcription, metabolism and development with the time of day [3–8]. In plants, environmental information including the light and temperature conditions is used to adjust the phase of the circadian oscillator, through the process of entrainment, so that the phase is aligned with the 24 h environmental cycle. This alignment between the circadian oscillator and the 24 h environmental cycle contributes to the fitness of plants [5].

Under natural conditions, environmental cues are a major determinant of temporal programs of gene expression [9]. For example, 97% of diel transcript profiles in field-grown rice can be predicted from meteorological data [9], and temperature cues regulate the alternative splicing of transcripts encoding circadian oscillator components in field-grown sugarcane [10]. Recent studies have provided insights into the diel organization of the transcriptome and metabolism, under field conditions, for several crops and *Arabidopsis* species [9–18]. A question that arises from these studies is how information from the circadian clock and environmental cues are processed to form integrated outputs. The diel dynamics of environmental signalling pathways, with defined inputs and outputs, are less well understood under natural conditions. The diel regulation of processes under controlled laboratory conditions does not necessarily reflect the dynamics in the field, even when using sophisticated replication of field conditions [15, 17]. Therefore, analysing the dynamics of cellular processes within complex fluctuating natural conditions allows us to elucidate how molecular mechanisms identified under controlled laboratory conditions operate in a natural setting [15]. Understanding this is valuable for translating laboratory studies into crop improvement. Furthermore, considering that plants comprise the major part of organismal biomass on Earth, it could contribute to forecasting ecosystem responses to increasingly unpredictable climates [19, 20].

To study the integration and transduction of circadian and environmental signals under naturally fluctuating conditions, we selected a well-characterized environmental signalling pathway as a model system. This model comprises the regulation of three components; a nuclear-encoded key component of the *Arabidopsis* circadian clock (*CIRCADIAN CLOCK ASSOCIATED 1*, *CCA1*), a nuclear-encoded regulator of chloroplast transcription (*SIGMA FACTOR 5*, *SIG5*), and the chloroplast-encoded gene *psbD*, which encodes the D2 protein of Photosystem II [21–23] (Fig. 1A). *CCA1* transcript abundance can be used as a proxy for the status of the circadian clock. *SIG5* is regulated closely by the circadian oscillator under constant conditions [22], and its transcript abundance responds to light, temperature and other environmental cues [21–33], which made us reason that SIG5 is an integrator of circadian and environmental information. SIG5 is imported into chloroplasts, and regulates transcription from the blue light responsive promoter of *psbD* (*psbD* BLRP) [21] (Fig. 1A). Thus, we assumed that SIG5-mediated signalling to chloroplasts involves a hierarchically-organized pathway, whereby *AhgCCA1* is positioned upstream from the regulation of *AhgSIG5* transcript accumulation, and *AhgpsbD* BLRP is positioned downstream of *AhgSIG5* activity (Fig. 1A). We also assumed that environmental signals might influence each component independently (Fig. 1A) [34]. We chose this pathway because it includes three major components representing distinct regulatory points of signal transduction, and has a relatively low level of complexity to evaluate circadian and environmental signal integration and transduction under realistic field conditions.

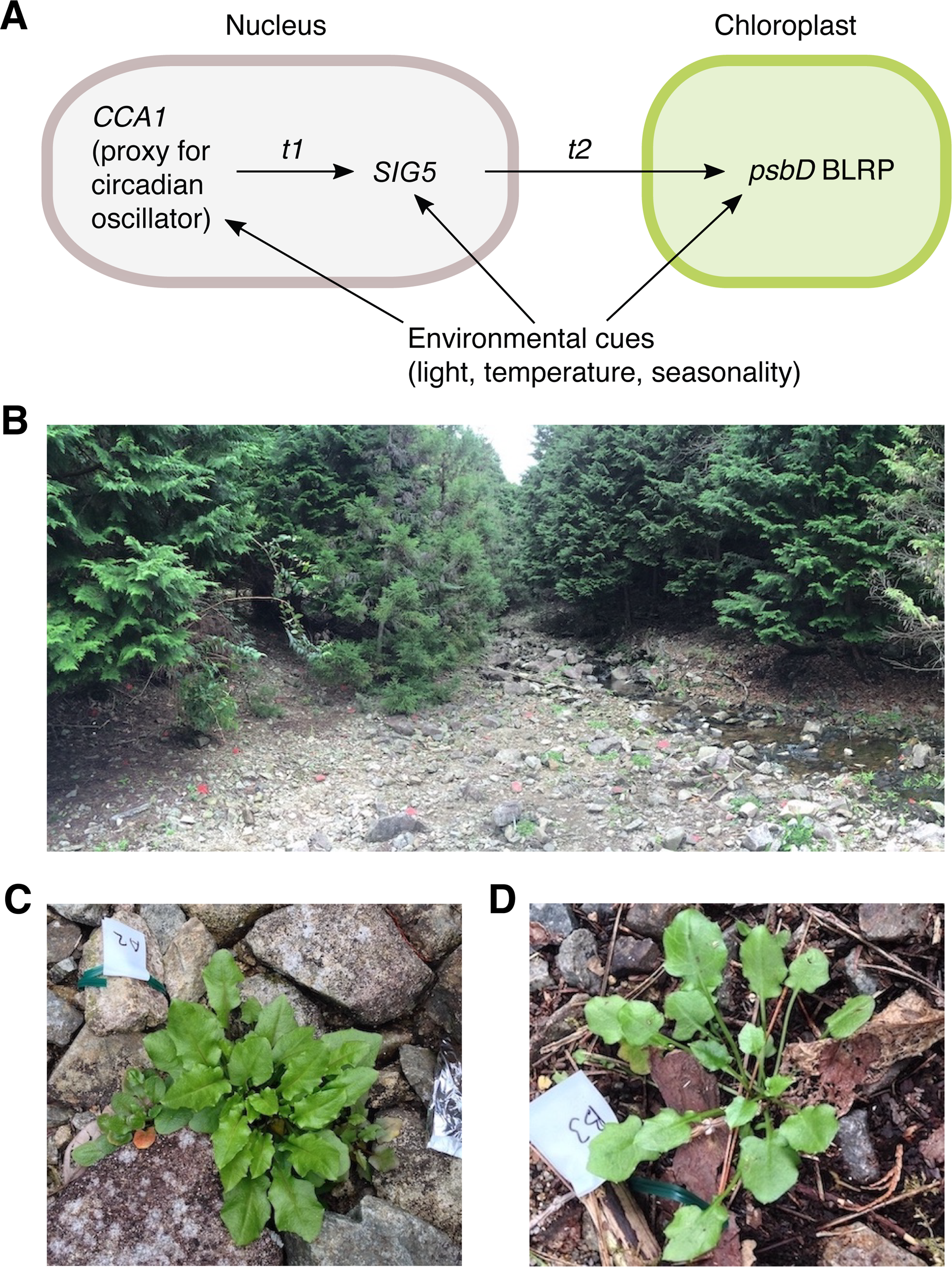

By targeting this model pathway, we combined time-series gene expression data with statistical modelling to investigate the temporal dynamics of signal integration and transduction in a fluctuating natural environment. We conducted the study using a natural population of a perennial *Arabidopsis* species (*Arabidopsis halleri* subsp. *gemmifera*, referred to as *A. halleri*) (Fig. 1B-D) [35]. The close-relatedness of *A. halleri* and *A. thaliana* makes it possible to identify pairs of homologous genes based on the sequence similarity (indicated by *Ahg* or *At* prefixes to gene names) [36]. The circadian clock-SIG5-*psbD* BLRP pathway is present in *A. thaliana* and *A. halleri*, and well-conserved across the vascular plants [37, 38]. We obtained several time series, during two seasons of the year, that monitored pathway function under representative light and temperature conditions in *A. halleri* in its natural habitat.

Interpreting time-series transcript data derived from complex fluctuating environments is a challenging task. We used statistical models, rather than models of biochemical kinetics [39], because this provides an effective tool for interpreting the influences of external factors on the response variables. We applied state-space modelling that integrates observations and the latent true state of a dynamic system into a statistical model. It is often used to interpret trends and oscillations in a system, and to evaluate the effects of external drivers on time-series data [40–42]. It explicitly formulates the system noise and observation error as state and observation equations, respectively. By incorporating an autoregressive trend and exogenous influences into a state space model, we estimated the dynamics of each pathway component that arises from its internal dynamics and external factors. Comparable approaches have allowed the investigation of diel and seasonal changes of transcriptome dynamics in *A. halleri* [12, 43, 44] and rice [9].

We evaluated evidence for causal regulation between the pathway components using convergent cross mapping (CCM), which provides a quantitative test for causation in time-series of coupled dynamical systems [45]. In contrast to our predictive state-space models, CCM evaluates whether the historical fluctuation in one variable can be used to estimate the fluctuation in a second variable [45]. This has been used previously to investigate a variety of ecological processes, including the seasonal regulation of gene expression by epigenetic marks in a natural *Arabidopsis* population [44] and the seasonal transcriptome in Japanese beech [46]. Using these approaches, we identified key roles for temperature and the circadian clock in the regulation of this signalling pathway under natural conditions. To our knowledge, this is the first investigation of circadian signal transduction in plants in their natural habitat, and the first use of these analytical approaches to understand circadian clock-regulated pathway in the field. Our approaches could be applicable to the study of many circadian-regulated processes under naturally fluctuating conditions.

## Results

### Biological data underlying models of signal transduction

Under controlled conditions of constant light, *AtCCA1* and *AtSIG5* transcript abundance are very well correlated (Fig. S1A-D; data from [3, 4, 6, 22, 47]). This correlation between *AtCCA1* and *AtSIG5* transcript abundance is absent under light/dark cycles (Fig. S1E, F; data from [22, 48]), suggesting that the integration of light and dark cues alters the diel regulation of *AtSIG5* transcript accumulation. To investigate these processes of signal integration under natural conditions, we acquired time-series of transcript abundance during spring (March) and autumn/fall (September), close to the spring or autumn equinox (Fig. S2; Fig. S3). Although both the spring and autumn equinoxes share 12 h photoperiods, they provide contrasting temperature regimes (cool and warm, respectively) (Fig. 2A, B; Fig. S4A, B), with irradiance levels determined by weather conditions (Fig. 2C, D, Fig. S4C, D). This allowed us to investigate temperature, light and seasonal influences upon SIG5-mediated signalling to chloroplasts, because this pathway is known to be affected by light and temperature in *A. thaliana* [21, 23, 24, 28, 33, 37]. We obtained data from areas with open sky and with vegetational shade, to include within our models the transcriptional responses to a wider range of irradiance levels (Fig. 2C, D; Fig. S4C, D). The “sun” and “shade” sampling sites were chosen by measurement of the ratio of red to far-red light (R:FR) (Fig. S5) and availability of plant patches, because *A. halleri* does not grow in deep shade. The total light intensity at the sun sampling site was 5 to 10-fold greater during March 2015 than during September 2015, depending on the time of day, due to weather differences (Fig. 2C, D; Fig. S4C, D). During March 2015, the study site temperature ranged from 0 °C to 14 °C at the sun site, and from 0 °C to 13 °C at the shade site (Fig. 2A, B; Fig. S4A). The temperature was always above 17 °C during September 2015, with greater diel fluctuations at the sun site (Fig. 2A, B; Fig. S4B).

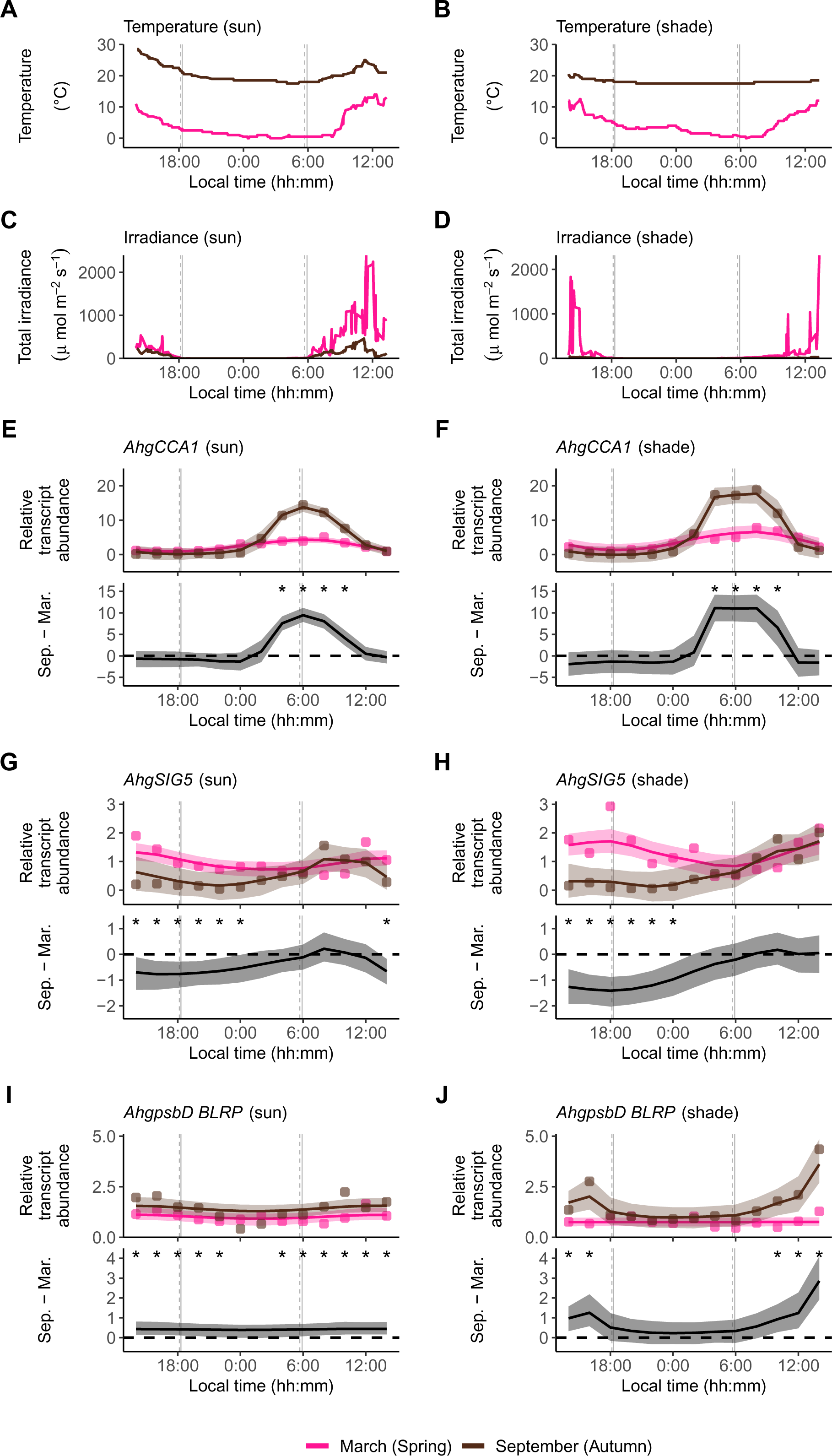

We monitored a 24 h cycle of abundance of *AhgCCA1*, *AhgSIG5* and *AhgpsbD* BLRP transcripts under each of these conditions, using RT-qPCR (Fig. S2; Fig. S3). Data from all the experiments were normalized to a single reference sample collected at midday, to enable comparability between sampling seasons and conditions. *AhgCCA1* oscillated with a 24 h cycle, reaching peak abundance at or just after astronomical dawn (Fig. S2E, F), whereas *AhgSIG5* and *AhgpsbD* BLRP reached peak abundance between the middle and end of the photoperiod, depending on conditions (Fig. S2G-J). The magnitude of oscillation of each transcript varied according to the sampling season and light conditions. *AhgCCA1* transcripts underwent a 5-(sun) and 7-fold (shade) change over the 24 h cycle during the March sampling, compared with a 98-fold (sun) and 190-fold (shade) change during the September sampling (fold-changes based on peak and trough abundance). This greater magnitude of oscillation occurred also for *AhgSIG5* during September (sun 38-fold; shade 33-fold) compared with March (sun 2.7-fold, shade 5.0-fold), and also for *AhgpsbD* BLRP during September (4.3-fold in sun and shade conditions) compared with March (sun 1.7-fold, shade 1.3-fold). Therefore, the relative magnitude of *AhgCCA1* transcript oscillation aligned well with that of *AhgSIG5* and *AhgpsbD* BLRP, whereby stronger diel cycling was present in September than in March, as suggested by the greater amplitude of *AhgCCA1* transcript during September sampling period compared with March. Furthermore, the magnitude (fold-change) of the 24 h oscillation of *AhgCCA1* was always greater than for *AhgSIG5,* and even less for *AhgpsbD* BLRP under any given condition. Whilst *AhgCCA1* underwent a greater daily fold-change under shade conditions compared with sun conditions, this was not reflected equivalently for *AhgSIG5* and *AhgpsbD* BLRP.

We compared the pathway dynamics between the spring and autumn sampling periods by using a smooth trend model (referred to here as STM). This is a type of state-space modelling that allows inference of a trend over time, where there is sampling noise and environmental stochasticity [49]. We estimated the parameters of a smooth trend model by Bayesian inference using the Markov Chain Monte Carlo (MCMC) approach [42, 50] to visualize the differences in transcript abundance between the spring and autumn sampling periods. Using this approach, we determined that the greater morning peaks of *AhgCCA1* expression in autumn compared to those in spring was statistically significant under both light conditions tested (the shaded areas of the lower panels of Fig. 2E, F represent the 95% confidence interval of the difference between seasons not containing zero around dawn). During both sampling seasons, *AhgSIG5* transcripts reached peak abundance between the middle and end of the photoperiod (Fig. 2G, H). This differs from *AtSIG5* transcript accumulation in *A. thaliana* under square-wave light/dark cycles under controlled conditions, where *AtSIG5* transcript abundance peaks around dawn [22]. The peak of the diel fluctuation of *AhgSIG5* was significantly greater during the March sampling period than during the September sampling period (95% confidence interval; Fig. 2G, H). Transcripts encoding the SIG5 regulatory target *AhgpsbD* BLRP (Fig. 1A) had a diel fluctuation during the September sampling season that reached a much greater peak under shade conditions compared with sun conditions (Fig. 2I, J), whereas there was very little diel fluctuation of *AhgpsbD* BLRP during the March sampling season.

*AhgCCA1* transcript abundance was significantly greater under shade compared with sun conditions during the photoperiod, during both sampling seasons (95% confidence interval; Fig. S4E, F). We did not identify the diminished *AtCCA1* oscillation that occurs under constant light with a very low R:FR [51] or on the shaded western side of crop fields around dawn [16]. In comparison, *AhgSIG5* transcript abundance was not altered consistently between sun and shade conditions (95% confidence interval; Fig. S4G, H). This tendency was also not observed for *AhgpsbD* BLRP transcript levels (95% confidence interval; Fig. S4I, J).

### Dynamics of environmental regulation of the signalling pathway

We represented the behaviour of the pathway components using a local level model with exogenous variables (referred to here as LLMX), a type of state-space modelling [52]. This allows the inclusion of a circadian or diel trend within a regression model that predicts transcript levels from environmental variables and the upstream regulatory component (Fig. 1A). The output of the Bayesian estimation reproduced well the dynamics of the observed *AhgSIG5* transcript level (Fig. 3A, B), with convergence of MCMC sampling (Fig. S6). The model estimated a clear diel trend of *AhgSIG5* transcript level that reached the trough level at dawn and the peak level between the middle and end of the photoperiod (Fig. 3C). The model also estimated significant negative and positive effects of ambient temperature and *AhgCCA1* on *AhgSIG5* transcript abundance, respectively (the error bars representing the 95% confidence interval of regression coefficients not containing zero; Fig. 3D). There was no significant effect of irradiance upon the estimation of *AhgSIG5* transcript abundance (Fig. 3D). Under controlled laboratory conditions, a temperature reduction leads to an increase in *SIG5* transcript abundance [33]. Therefore, the estimated negative relationship between ambient temperature and *AhgSIG5* transcript levels (Fig. 3D) could suggest a similar regulation in *A. halleri* under naturally fluctuating conditions. If the temperature input to the model was set to a constant value (the mean temperature across all conditions) and the estimated parameter values in this model were used, the greater *AhgSIG5* transcript abundance during March disappeared, whereas the phase of the dynamics was not affected (compare Fig. S7A, B and Fig. S7C, D). This suggests that temperature is a key predictor of the difference in *AhgSIG5* transcript abundance between seasons. Setting the irradiance input to a constant level (the mean irradiance across all conditions) did not affect the *AhgSIG5* dynamics (compare Fig. S7A, B and Fig. S7E, F). Setting a constant *AhgCCA1* transcript level shifted the phase of *AhgSIG5* dynamics in September (compare Fig. S7A, B and Fig. S7G, H), suggesting that *AhgCCA1* input partially contributes to the prediction of the *AhgSIG5* oscillation.

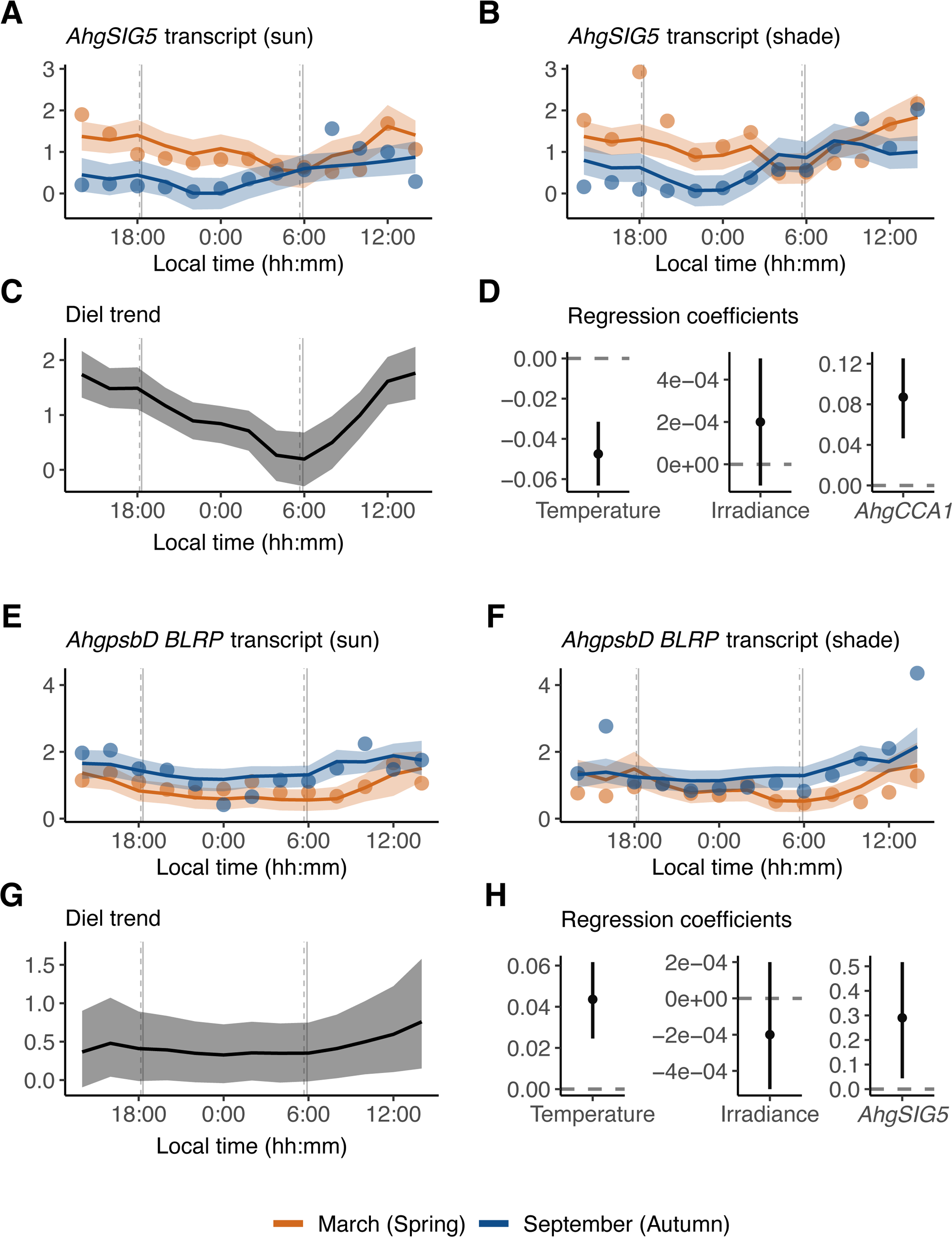

Diel fluctuations of *AhgpsbD* BLRP transcript abundance were reproduced reasonably well by the model (Fig. 3E, F), with convergence of MCMC sampling (Fig. S8). The model predicted a diel trend of *AhgpsbD* BLRP transcript level that increased towards the end of the photoperiod (Fig. 3G). The model estimated a significant positive effect of ambient temperature on *AhgpsbD* BLRP transcript abundance (95% confidence interval; Fig. 3H). In *A. thaliana* under controlled conditions, *psbD* BLRP transcript accumulation is regulated by SIG5 [21], and within our model, there was also a significant positive coefficient of regression between *AhgSIG5* and *AhgpsbD* BLRP transcript levels (95% confidence interval; Fig. 3H). Setting the temperature input as a constant value (the mean temperature among all conditions) with estimated parameter values in this model, greater *AhgpsbD* BLRP transcript abundance in September disappeared, and the diel dynamics became slightly flatter (compare Fig. S9A, B and Fig. S9C, D). This suggests that temperature is a key predictor of the difference in *AhgpsbD* BLRP transcript abundance between seasons. Constant irradiance did not affect the *AhgpsbD* BLRP dynamics (compare Fig. S9A, B and Fig. S9E, F). Constant *AhgSIG5* transcript level caused apparent flatter *AhgpsbD* BLRP dynamics (compare Fig. S9A, B and Fig. S9G, H), suggesting that *AhgSIG5* input makes a partial prediction of the *AhgpsbD* BLRP oscillation.

Overall, this analysis identifies that the ambient temperature, rather than the irradiance, was important for predicting the seasonal differences in the transcript abundance of *AhgSIG5* and *AhgpsbD* BLRP under naturally fluctuating conditions. This contrasts evidence from *A. thaliana* under controlled conditions, where light inputs have a key role [21, 23]. In addition, the circadian clock (*AhgCCA1*) affected the diel oscillation of *AhgSIG5* transcript abundance (Fig. 3D; Fig. S8G, H), and *AhgSIG5* affected the diel oscillation of *AhgpsbD* BLRP transcript abundance (Fig. 3H; Fig. S9G, H). More specifically, seasonal difference in temperature affected the base levels of *AhgSIG5* and *AhgpsbD* BLRP transcripts, while the diel oscillation patterns of *AhgSIG5* and *AhgpsbD* BLRP transcripts were affected by the transcript levels of their upstream genes rather than diel temperature fluctuations.

The significant effects of temperature on *AhgSIG5* and *AhgpsbD* BLRP transcripts (Fig. 3D, H), and the restriction to specific periods of the day of significant differences between the transcript dynamics under the different temperature conditions of March and September (Fig. 2E-J), is reminiscent of the concept of circadian gating. Circadian gating is the process whereby the circadian clock restricts certain biological processes to specific times in the 24 h cycle [53]. In plants, this can take the form of a circadian rhythm in the magnitude of the response to identical environmental stimuli given at different times of day [8].

### Temporal gating of temperature regulation of SIG5-mediated signalling to chloroplasts under natural conditions

Our LLMX analysis suggests that under natural conditions, lower ambient temperatures might upregulate *AhgSIG5* transcript levels, and greater ambient temperatures might upregulate *AhgpsbD* BLRP transcript levels (Fig. 3D, H). To test this hypothesis and further examine whether a circadian gating-like process might occur, we applied moderate temperature manipulations to adjacent patches of *A. halleri* plants, in the field, using custom-designed equipment (Fig. 4A; Fig. S10A-D). We collected 24-h time-series of RNA samples from these plant patches (Fig. S10E-G), and interpreted the data with smooth trend models (STM; Fig. 4B-D). This experiment occurred during September 2016, to take advantage of the greater magnitude of oscillation of the transcripts observed during experimentation during September 2015 compared with March 2015 (Fig. 2E-J). *AhgCCA1* transcripts underwent a substantial fold-change fluctuation over the 24 h cycle, ranging from 210-fold (moderate temperature reduction) to 339-fold (ambient temperature) and 1,230-fold (moderate temperature increase) (Fig. S10E). Interestingly, a moderate temperature reduction treatment during September 2016 did not decrease the fluctuation of *AhgCCA1* transcripts to a level equivalent to the March 2015 sampling season (5 to 7-fold fluctuation), suggesting that either there is a tipping point temperature below which *A. halleri CCA1* transcripts have a substantial loss of rhythmicity, or that the past temperature history of the plants during each season can influence the amplitude of oscillation of this circadian clock transcript.

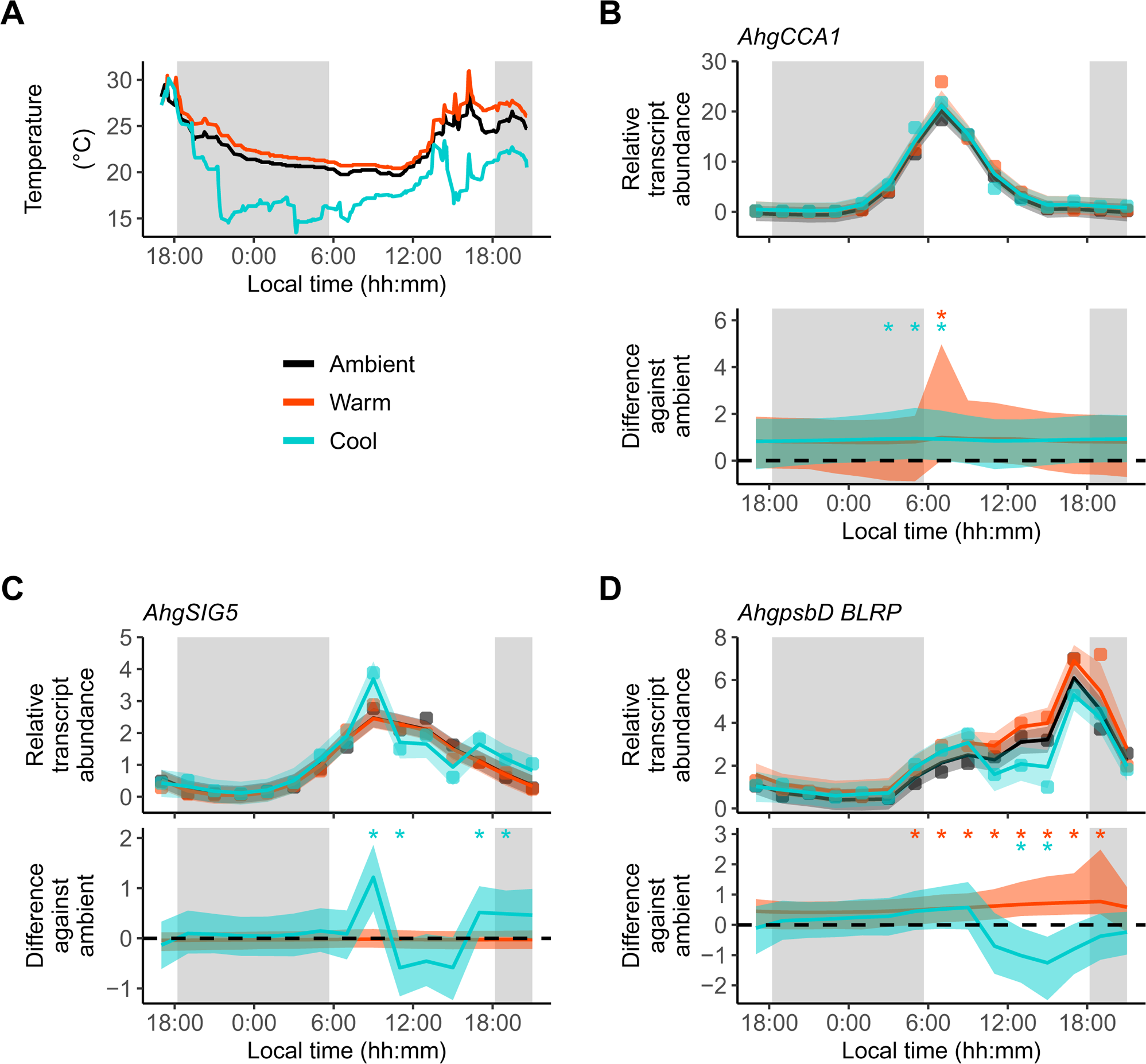

Both the moderate temperature increase and temperature reduction treatment caused a small significant upregulation of *AhgCCA1* transcript abundance around dawn relative to the ambient temperature control (significance at > 95% probability; Fig. 4B). The moderate temperature increase treatment was without effect upon *AhgSIG5* transcript abundance (Fig. 4C). In comparison, the temperature reduction treatment upregulated *AhgSIG5* transcripts significantly after dawn and around dusk, relative to the ambient temperature control (significance at > 95% probability; Fig. 4C). This is consistent with the negative coefficient of regression of temperature on *AhgSIG5* transcript under naturally fluctuating conditions (Fig. 3D), and with the upregulation of *A. thaliana SIG5* by a short cold treatment under laboratory conditions [21, 33]. The restriction of the response of *AhgSIG5* transcripts to the moderate temperature reduction to certain times of day (Fig. 4C) supports the notion that there is temporal gating of the response of *AhgSIG5* to temperature under naturally fluctuating conditions.

Transcripts for the chloroplast target of SIG5, *AhgpsbD* BLRP, were also altered by temperature manipulation (Fig. 4D). The moderate temperature elevation significantly increased *AhgpsbD* BLRP transcript levels relative to the control, whereas the moderate temperature reduction significantly reduced *AhgpsbD* BLRP transcripts relative to the control (significance at > 95% probability; Fig. 4D). This is consistent with the positive coefficient of regression of temperature on *AhgpsbD* BLRP transcript abundance under naturally fluctuating conditions (Fig. 3H). The response of *AhgpsbD* BLRP transcripts to temperature manipulations was restricted to the photoperiod, which might be because chloroplast DNA binding and transcription by PEP generally requires light [54–58].

### Evaluation of causal regulation within a circadian-regulated signalling pathway under natural conditions

We performed CCM analysis of the relationship between *AhgCCA1* and *AhgSIG5*, and of the relationship between *AhgSIG5* and *AhgpsbD* BLRP, using a variety of time-delays between each pair of variables. We considered potential time-delays in CCM analysis [59] because the abundance of each related transcript was monitored at each timepoint, but their responses to each other might not be instantaneous. For example, in *A. thaliana* under controlled square-wave light/dark cycle conditions, *AtCCA1* transcript abundance peaks at dawn, *AtSIG5* approximately 3 h after dawn, and *AtpsbD* BLRP approximately 6 hours after dawn [22].

To prepare for CCM analysis, we determined the optimal embedding dimension for each variable (*AhgCCA1*, *AhgSIG5* and *AhgpsbD* BLRP; Fig. S11). A feature of CCM analysis that can confuse readers is that the direction of successful prediction is *opposite* to that of causality (Fig. 5A). Subsequent analysis with CCM identified a significant prediction (cross map skill) of *AhgCCA1* transcript abundance from *AhgSIG5* transcript abundance (Fig. 5B) when no time delay was incorporated between the variables. Furthermore, a significant prediction of *AhgSIG5* from *AhgpsbD* BLRP transcript abundance occurred when no time delay was incorporated, and potentially also with a time delay between *AhgSIG5* and *AhgpsbD* BLRP (Fig. 5C). In both cases, the cross map skill exceeded a 95% interval of the diel surrogate, which imitates the same degree of oscillation of the explanatory variable, but where the variation is randomized (Fig. 5B, C). When the library size (number of time points used to reconstruct a state space) was increased, cross map skill was improved for each time lag, attaining a significant cross map skill (Fig. 5D-F), suggesting convergence (a key property that distinguishes causation from simple correlation) [45]. Overall, CCM analysis suggests that there is a causal relationship between fluctuations in *AhgCCA1* and *AhgSIG5*, and *AhgSIG5* and *AhgpsbD* BLRP, and that there is a potential time-delay from nuclear genome-encoding *AhgSIG5* to chloroplast genome-encoding *AhgpsbD* BLRP in a natural plant population.

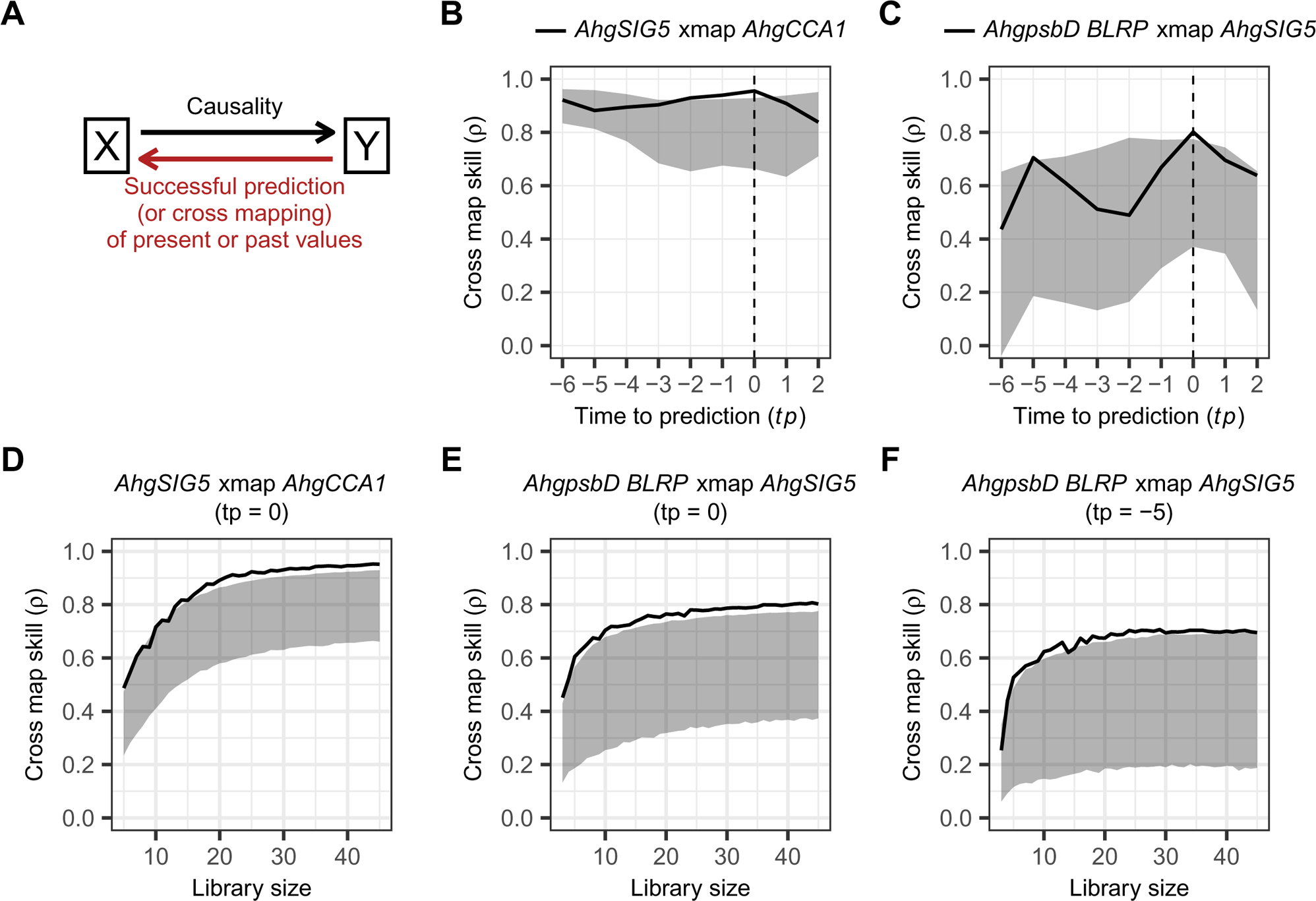

## Discussion

We deployed a set of approaches that allowed us to gain insights into the integration and transduction of circadian and environmental signals in a natural plant population. Using a modelling approach, we predicted that the circadian clock and ambient temperature are potential regulators of *AhgSIG5*-mediated signalling to chloroplasts, in a natural population of *A. halleri*. There was a significant influence of *AhgCCA1* on *AhgSIG5* transcript abundance, and of *AhgSIG5* on *AhgpsbD* BLRP transcript abundance (Fig. 3D, H). Furthermore, convergent cross mapping analysis predicted significant causal relationships within this signalling pathway in the field (Fig. 5). Therefore, it appears that in a natural plant population, a signal is communicated from the circadian oscillator (using *AhgCCA1* as a proxy) to the signalling pathway output of *AhgpsbD* BLRP. One interpretation is that the pathway couples the circadian oscillator and temperature response processes to chloroplast gene transcription under naturally fluctuating conditions.

We identified seasonal differences in the maximum accumulation of *AhgCCA1*, *AhgSIG5* and *AhgpsbD* BLRP. *AhgCCA1* and *AhgpsbD* BLRP had significantly lower peak accumulation during the spring sampling period compared with the autumn sampling season (Fig. 2E, F, I, J). In comparison, *AhgSIG5* had significantly greater peak accumulation during the spring compared with the autumn sampling season (Fig. 2G, H). The difference in *AhgCCA1* dynamics between these sampling seasons likely reflects the decreased amplitude of the circadian oscillator that occurs under lower temperature conditions, in both controlled laboratory environments and the field [12, 60–62]. The difference in the seasonal response of *AhgSIG5* compared with *AhgCCA1* suggests that an additional temperature input into this pathway occurs between the circadian oscillator and *AhgSIG5*. In *A. thaliana*, *AtSIG5* transcripts are upregulated by short cold temperature treatments [21, 33]. In our field experiment, *AhgSIG5* transcript accumulation showed a negative coefficient of regression with temperature (Fig. 3D), with its accumulation increased during the morning and around dusk by a low temperature manipulation (Fig. 4C). This suggests that under lower temperature conditions, *AhgSIG5* transcript abundance will increase. Therefore, the lower temperatures of the spring sampling season compared with the autumn sampling season (Fig. 2A, B) might explain the significantly greater abundance of *AhgSIG5* transcripts at many times of day during the spring. A further feature of interest within the data is the relative robustness of the dawn timing of peak *AhgCCA1* abundance, irrespective of the environmental conditions (Fig. 2E, F), which is consistent with a previous field study [12]. In *Arabidopsis*, *AtCCA1* is light-induced [63, 64], yet under field conditions *AhgCCA1* reached a peak around astronomical dawn when the irradiance was relatively low, and was unresponsive to increased light intensity later in the day (Fig. 2C-F). Furthermore, its peak time was relatively unaffected by seasonal differences in light conditions (Fig. 2C-F). This reveals an astonishing consistency of the phase of expression of a clock gene in plants under complex fluctuating environments, which might be accomplished through the complexity of the *Arabidopsis* circadian oscillator that is thought to confer robustness to its regulation [65, 66], and perhaps a restriction of light-induced phase shifts to certain times of day due to the gating of light inputs. Mathematical modelling indicates that the interplay between different simultaneous circadian entrainment cues might reinforce circadian clocks [67], although the timing of individual cues applied in combination can also cause changes in the phase [68]. The processes that align the phase of the plant circadian clock in plants with growing environments, under naturally fluctuating conditions, represent a fascinating topic for future investigation.

Because we considered *AhgpsbD* BLRP to represent the ultimate output from the signalling pathway (Fig. 1A), our analysis suggests that environmental inputs occurred within at least three positions in the pathway; first, in the regulation of *AhgCCA1* transcript accumulation by the season or temperature, second, in the regulation of *AhgSIG5* transcript accumulation by temperature, and third, in the regulation of *AhgpsbD* BLRP transcript accumulation by temperature. These environmental inputs might occur through several biologically independent processes. One possibility is temperature inputs to the circadian clock mediated by temperature-responsive components, such as the evening complex [69, 70]. A further mechanism could be the regulation of *AhgSIG5* by HY5, which is a known regulator of SIG5 that participates in low-temperature gene regulation [21, 27, 71] and underlies a response of *AtSIG5* transcripts to cold in *A. thaliana* [33]. Furthermore, there might be direct effects of light upon sigma factor activity in chloroplasts through, for example, redox regulation [72] or light- and temperature-regulation of chloroplast protein import [73]. We did not consider here the short- or long-term history of light or temperature conditions upon leaves prior to experimentation. However, the past temperature conditions can affect *AhgCCA1* oscillations, since the sensitivity of *AhgCCA1* oscillations to a moderate temperature reduction treatment during September 2016 was limited compared with its response to the lower ambient temperature conditions during Spring 2015. Other biotic or abiotic factors such as water availability, relative humidity, and atmospheric CO_2_ concentration might also have effects that we did not include in our models.

Statistical modelling of the transcriptome of field-grown *Oryza sativa* (rice) concluded that the main environmental driver of *OsSIG5* (*Os05g0586600*) transcript accumulation is temperature [9]. In this case, the temperature had a negative regression coefficient with *OsSIG5* [9]. This is consistent with our finding of a negative coefficient of regression of temperature on *AhgSIG5*. The study of the rice transcriptome in the field [9] did not monitor chloroplast-encoded transcripts, so a direct comparison between *AhgpsbD* BLRP and our data is not possible.

A key finding from our work is the detection of a 24 h fluctuation of the influence of the temperature upon each of the three genes (Fig. 4). One interpretation is that there is a diel cycle of sensitivity of these pathway components to temperature cues, with their response to temperature restricted to certain times of day, reminiscent of the process of circadian gating [53]. Our findings from the field are corroborated by laboratory experiments, which demonstrate that the circadian oscillator gates its own response to temperature [74], and the response of *AtSIG5* to blue light and cold temperatures is gated by the circadian oscillator [22, 33]. Statistical modelling approaches applied to diel cycles of the rice transcriptome in the field have identified 24 h oscillations in the coefficient that describes the relationship between temperature and transcript levels, which is also reminiscent of circadian gating [9]. In field experiments, it is difficult to link the fluctuating temperature sensitivity to the circadian oscillator causatively, although a combination of field and laboratory experiments investigating the environmental regulation of *FLOWERING LOCUS T* (*FT*) expression suggest that the circadian clock can gate light quality and temperature responses of *FT* under field conditions [18, 75]. Taken together, these findings are important because they suggest that processes leading to circadian gating might operate in natural plant populations to modulate their environmental responses.

Our investigation provides insights into molecular aspects of signal transduction in plants under field conditions. This represents a relatively under-studied topic, and we adopted quantitative approaches to interpret relatively noisy transcript data collected under complex fluctuating environments, to investigate the functioning of a specific pathway. This allowed us to infer multiple positions of environmental inputs into the pathway, and temporal gating of a response to temperature. The approaches used provide a framework to study environmental signal integration in all organisms under field conditions, providing opportunities to investigate whether and how mechanisms identified in the laboratory might function in the field. This could be important to understand rhythmic biological responses within an increasingly unpredictable climate.

## Materials and Methods

### Field site and plant material

Our experiments used a naturally-occurring population of *Arabidopsis halleri* subsp. *gemmifera* (Matsum.) O’Kane & Al-Shehbaz growing beside a forested stream in Hyogo Prefecture, Japan (Omoide-gawa site; 35°06’ N, 134°55’ E, elevation 190–230 m) [12, 43, 76] (Fig. 1B). We selected *A. halleri* as an experimental model for several reasons [35]. First, it has high nucleotide sequence identity and good synteny with *A. thaliana* [36]. Second, unlike *A. thaliana*, the perennial life history of *A. halleri* allows investigation of transcriptional responses across the seasons [2, 76]. Many individuals are clones because the species propagates by producing clonal rosettes as well as by seeds, which allows repeated sampling from single genotypes. Furthermore, *A. halleri* is metal tolerant and occurs in natural habitats that are relatively free from other vegetation due to contamination by heavy metals, which provides experimentally-convenient sites enriched with many *A. halleri* plants [76]. *Arabidopsis halleri* subsp. *gemmifera* at this site was previously identified by examination of museum and herbarium specimens, and a nearby population provided material for sequencing the *A. halleri* genome [36, 76].

*AhgCCA1* (*g097040*) and *AhgSIG5* (*g25274*) were identified from *A. halleri* genome Version Ahal2.2 [36]. *AhgCCA1* has 94.8% coding sequence identity and 93.3% protein sequence identity with *AtCCA1*. *AhgSIG5* has 94.9% coding sequence identity and 95.0% protein sequence identity with *AtSIG5*. Chloroplast-encoded *psbD* is not annotated within Version Ahal2.2 of the *A. halleri* genome, and we identified this instead within scaffold 2 of an *A. halleri* reference transcriptome [77]. The *AhgpsbD* BLRP promoter region, which was our focus, has a 100% sequence identity with *AtpsbD* BLRP [55].

### Experimental conditions for field sampling of A. halleri

The first sets of samples were obtained under natural conditions without temperature manipulation during 24 – 26 March 2015 and 15 – 17 September 2015, which were close to the spring and autumn equinox at the field site, respectively. We exploited variations in environmental conditions across the field site, and sampled leaves from the locations nominated as “sun” and “shade” sites on successive days. At “sun” locations, plants received direct sunlight during the day, and at “shade” locations plants received sunlight filtered by surrounding vegetation for most of the day with the sites identified by measurement of the ratio of red to far red light (Fig. S5; R:FR calculated as the photon irradiance from 660 to 670nm divided by the photon irradiance from 725 to 735nm [78]). During March 2015, plants received more direct sunlight, whereas during September 2015 the light was scattered through sky overcast with clouds.

The second sets of samples were obtained under natural conditions with manipulation of the temperature conditions around patches of plants during 13 – 14 September 2016, which was close to the autumn equinox at the field site. In addition to control plants that were not manipulated (Fig. S10A), we applied two temperature treatments. These were (1) a continuous temperature increase (Fig. S10B), whereby plants were covered with clear plastic horticultural domes to block air currents and trap warm air; (2) a continuous temperature reduction, using a custom device that passed air through a duct within a heat-exchanging ice-filled polystyrene box and expelled the chilled air into a clear horticultural dome covering the plants, with chilling augmented by small ice packs within the dome (Fig. S10C).

### Field sampling for transcript analysis

Across all experimental conditions, the same sampling and RNA isolation procedures were used. At 2 h intervals, a fully expanded rosette leaf was excised with dissecting scissors from 6 replicate plants for each condition. The time-courses using naturally occurring sun and shade conditions each comprised 13 sampling timepoints over a total of 26 hours (from 14:00 on the first day to 16:00 on the second day). The time-courses involving artificial temperature manipulations comprised 15 sampling timepoints over a total of 30 hours (from 17:00 on the first day to 21:00 on the second day). The same replicate plants were sampled repeatedly through each time-series, but different plant patches were used in different sampling seasons. Sampled leaves were placed immediately into individual microtubes containing at least 400 µL RNA*later* Stabilization Solution (Thermo Fisher Scientific, Waltham, MA, USA). Scissors and forceps were cleaned with 70% (w/v) ethanol between samples. After sampling, tubes were placed temporarily on dry ice for up to 2 hours, at −40 °C for 3 days in a portable freezer during transfer to the laboratory, and then at −80 °C until RNA isolation. During hours of darkness, sampling occurred using green-filtered head torches.

We wished to ensure that the abundance of transcripts could be compared between each sampling season. We normalized all transcript measurements to the transcript levels in one sample. Therefore, we obtained this reference sample for normalization of all RT-qPCR experiments in the study by pooling RNA from 10 leaves sampled at noon on 26 March 2015, from 10 healthy plants across the field site that were each separated by at least 1 metre. This provided a reference cDNA sample against which all RT-qPCR analyses from all sampling seasons were normalized within the ΔΔCt method [79], to allow comparability between all datasets. The reference sample was collected at midday because all transcripts under investigation were expressed to some extent at that time point. In all experiments, dawn and dusk were defined as the astronomical (solar) time of sunrise and sunset.

### RNA isolation and RT-qPCR

Frozen samples containing RNA*later* were defrosted in a cold room for 4 hours, the RNA*later* was removed, and leaf tissue was transferred to new dry tubes and frozen in liquid nitrogen. Frozen tissue was ground with a TissueLyzer (Qiagen, Hilden, Germany) and total RNA was isolated from the powdered plant material using Macherey-Nagel Nucleospin II RNA extraction kits (Thermo Fisher Scientific). RNA concentrations were determined using a Nanodrop spectrophotometer (Thermo Fisher Scientific). cDNA was synthesized using a High Capacity cDNA Reverse Transcription Kit (Thermo Fisher Scientific) and random primers supplemented with RNAase inhibitor (Thermo Fisher Scientific), as described previously [22, 23]. 1:500 cDNA dilutions were analyzed using Brilliant III Ultra-Fast SYBR Green QPCR Master Mix (Agilent Technologies, Santa Clara, CA, USA), required primer pairs (Table S2), and Agilent Mx3005P qPCR instrument. Primers were designed using the PrimerQuest™ Tool (Integrated DNA Technologies, Coralville, IA, USA). Results were normalized using the ΔΔCt method to *AhgACTIN2* [22, 23]. *AhgACTIN2* is encoded in *A. halleri* by locus *g21632* [36] and has 97.8% coding sequence identity with *A. thaliana ACTIN2* (*At3g18780*). We selected *AhgACTIN2* as a reference transcript, because it has been used previously as a reference transcript for experiments involving *A. halleri* at this field site [43], *AhgACTIN2* does not fluctuate across the seasonal cycle [12], and in our experiments, the RT-qPCR Ct for *AhgACTIN2* did not oscillate across the diel cycle (*AhgACTIN2* time-series arrhythmic after analysis with JTK_CYCLE test for rhythmicity [80]; Fig. S12).

### Environmental monitoring

The temperature and irradiance were measured beside the plants during sampling. The temperature at each location, for each temperature manipulation, was monitored with EL-USB-2 data loggers (Lascar Electronics, Whiteparish, UK) at 5-minute intervals.

Temperature loggers were wrapped in aluminium foil to prevent surface heating by solar radiation. Irradiance was measured using a CC-3-UV-S cosine corrector connected to a USB2000+ spectrometer with a QP400-2-UV-VIS fibre optic cable (Ocean Optics, Dunedin, FL, USA). Ambient light spectra (200 nm to 900 nm) were collected every 5 minutes over the 14 hours of light during each day of sampling using OceanView software (Ocean Optics) on a laptop PC, controlled by a custom script. The spectrometer and computer were powered using portable lithium battery packs (Powertraveller, Hampshire, UK).

### Smooth trend model

The smooth trend model (STM) allows inference of a trend within time-series data that contains both sampling noise and biological stochasticity, and also allows a level of statistical confidence to be applied to that trend. The STM to assess the difference in transcript abundance between March and September under sun and shade conditions (Fig. 2) was defined by the equations:

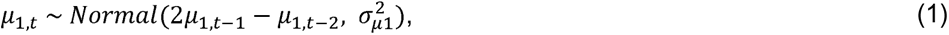

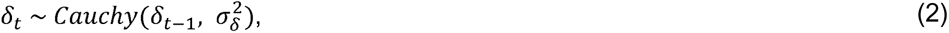

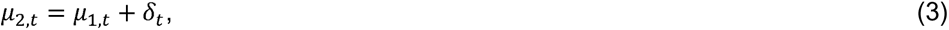

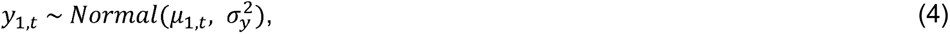

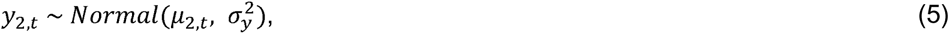

where *µ*_l_ and *µ*_2_ are the smooth trend components in March and September in 2015, respectively, *δ* is the time-varying difference between the two seasons, and *y*_1_ and *y*_2_ are the observed transcript abundance in the two seasons. *t* = (1, 2,…, 13) is the time point at two-hour intervals. The same STM was used to analyze the difference in transcript abundance between sun and shade conditions in March and September (Fig. S4).

For the models of the three (ambient, warm and cool) conditions in the temperature manipulation experiment (Fig. 4), additional *δ*, *µ* and *y* were considered:

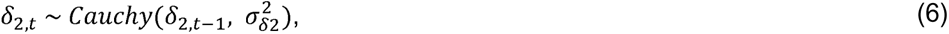

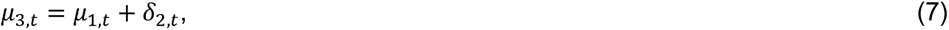

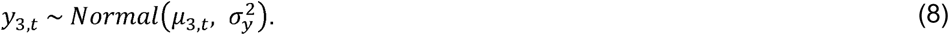

The parameters of the models were estimated by Bayesian inference using the Markov Chain Monte Carlo (MCMC) approach. We note that unlike classical hypothesis testing methods, multiple comparisons do not raise a problem in a Bayesian multilevel modelling [81]. The statistical models were written in the Stan language and the programs were compiled using CmdStan (v2.24). To operate CmdStan, the cmdstanr package (v0.4.0) in R was used. After 1,000 warm-up steps, 1,000 MCMC samples were obtained by thinning out 3,000 MCMC samples for each of four parallel chains. Thus, 4,000 MCMC samples were obtained in total. As prior distributions, we used a positive uniform distribution for σ, cauchy distribution for *δ* (equations 2 and 6), and a normal distribution for *µ* and *y* (equations 1, 4, 5 and 8). A cauchy distribution of the priors was used to represent the difference between two time series (*δ*) because this distribution corrects better for the influence of outliers, due to its relatively long tails and its efficiency for detecting change points in time series data [82, 83].

### Local level model with exogenous variables

The local level model with exogenous variables (LLMX) to analyze a diel trend and the effect of environmental variables on transcript abundance (Fig. 3) was defined by the equations:

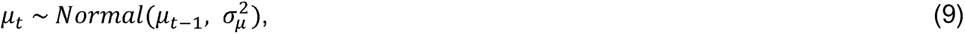

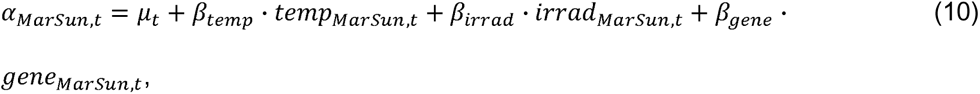

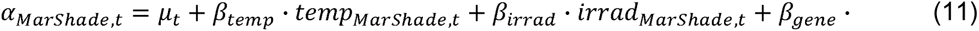

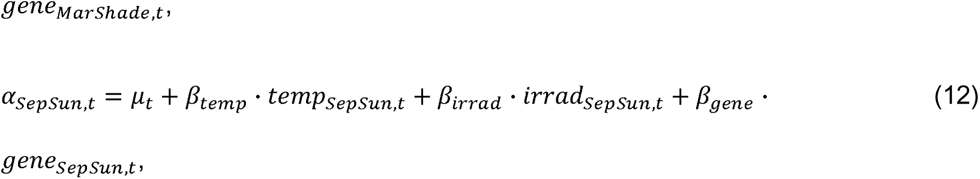

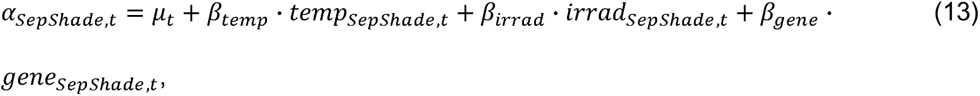

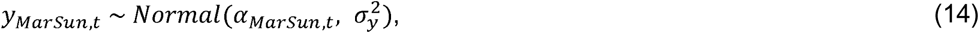

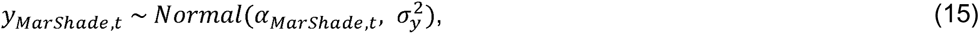

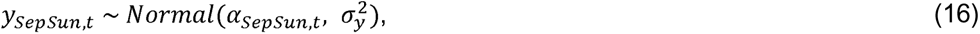

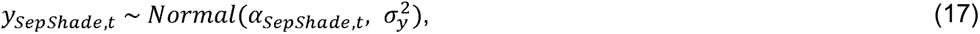

where *µ* is the autoregressive trend component that is common among conditions, *β* is the regression coefficient, α is the true state of transcript abundance, *y* is the observed transcript abundance, and σ^2^ is the variance. The subscripts, *temp, irrad, gene, Mar, Sep, Sun* and *Shade* represent temperature, irradiance, an upstream gene, March, September, sun condition and shade condition, respectively. The mean transcript abundance of the upstream gene (i.e., *AhgCCA1* in the *AhgSIG5* model and *AhgSIG5* in the *AhgpsbD* BLRP model) was used as one of the inputs. *t* = (1, 2,…, 13) is the time point at two-hour intervals.

The parameters of the models were estimated by Bayesian inference using the Markov Chain Monte Carlo (MCMC) approach (Table S2, S3). The statistical models were written in the Stan language and the programs were compiled using CmdStan (v2.24). To operate CmdStan, the cmdstanr package (v0.4.0) in R was used. After 3,000 warm-up steps, 1,000 MCMC samples were obtained for each of the four parallel chains, and thus 4,000 MCMC samples were obtained in total. As prior distributions, we used a uniform distribution for *β*, a positive uniform distribution for a, and a normal distribution for *µ* and *y* (equations 9 and 14-17).

### Embedding dimension

In the empirical dynamic modelling (EDM), time series is embedded into time-lagged series, a procedure known as state space reconstruction [84, 85]. The embedding dimension *E* is the dimension (i.e. the number of time-lagged series) used to reconstruct the state space. Prior to convergent cross mapping (CCM), the optimal *E* value was determined for each of *AhgCCA1*, *AhgSIG5* and *AhgpsbD* BLRP in temperature manipulation experiments in September 2016, by univariate simplex projection [86], using the simplex function of the rEDM package (v0.7.5) of R. We determined the optimal *E* values showing the maximum forecast skill ρ (Fig. S11A). The optimal *E* value of 8 for *AhgCCA1* was relatively large, considering those of *AhgSIG5* and *AhgpsbD* BLRP, and data points available in this analysis. We adopted *E* value of 2 as the optimal value for *AhgCCA1*, because ρ is higher than 0.9 for *E* values from 2 to 8 (Fig. S11A) and the model prediction well fits the observed data at *E* value of 2 (Fig. S11B). The optimal *E* values determined were 2 for *AhgCCA1*, 4 for *AhgSIG5*, and 2 for *AhgpsbD* BLRP. The model predictions well fit the observed data for these genes at the determined *E* values (Fig. S11B-D).

### Convergent cross mapping

When a causal relationship exists from a variable *X* to another variable *Y*, the information of *X* can be found in the time series of *Y*, meaning that the prediction of *X* is possible using the information of *Y*. In convergent cross mapping (CCM) based on simplex projection, one predicts the nearest neighbors of *X* at time *t* in the reconstructed lagged trajectory, using their time-corresponding points of *Y* [45]. When this prediction (called cross mapping) is successful, the causality from *X* to *Y* is assumed. The cross map skill (i.e. prediction skill) is evaluated by Pearson’s correlation coefficient (ρ) between predicted and observed values. We performed CCM between *AhgCCA1* (*X*) and *AhgSIG5* (*Y*), and between *AhgSIG5* (*X*) and *AhgpsbD* BLRP (*Y*) for the temperature manipulation experiments (September 2016), using the ccm function in the rEDM package (v0.7.5) of R. We considered time delay in the interactions by changing the parameter tp from −6 to 2 (negative, past; positive, future) in the ccm function. When causality exists, optimal predictability is expected to occur for tp ≤ 0, i.e, prediction of past values of *X* from *Y* [59]. We used a technique known as multispatial CCM [87], to utilize the relatively small number of time points of 15/condition. Since we used three temperature conditions (ambient, warm and cool), we applied the total of 45 time points to CCM. To test the significance of cross map skill, we produced 1,000 diel surrogate time series for *X* into which a similar level of oscillation was incorporated, but the sequence of data deviation from the oscillation was randomized. We used three criteria for significant causality. First, optimal predictability occurs for tp ≤ 0, second, cross map skill is above the 95 % interval of diel surrogates at the optimal time lag (tp), third, cross map skill is improved according to the increase in a library size (number of time points used to reconstruct a state space), known as convergence [45].

## Acknowledgements

We thank Noriane M. L. Simon and Tasuku Ito for experimental assistance, and Paige E. Panter and Deirdre A. Lynch for constructive feedback. This research was funded by BBSRC (UK) (BB/I005811/2; BB/J014400/1; Institute Strategic Programme GEN BB/P013511/1; Institute Strategic Programme bRiC BB/X01102X/1), The Royal Society (IE140501), The Leverhulme Trust (RPG-2018-216), the Japan Society for Promotion of Science (JSPS KAKENHI, JP21H04977, JP21H05659, JP21K15164), the Japan Science and Technology Agency (JST, CREST no. JPMJCR1501) and the Bristol Centre for Agricultural Innovation. DLCR is grateful to Consejo Nacional de Ciencia y Tecnología (Mexico) for awarding a scholarship. This research was conducted through the Joint Usage programme of the Center for Ecological Research of Kyoto University.

## Author contributions

DLCR, HN, JS, MNH, HK and AND designed and performed experimentation; HN, DLCR, TM, LLBD, HK and AND analyzed and interpreted data, and DLCR, HN, LLBD, HK and AND wrote the paper.

## Data and code availability

Source data are provided in Spreadsheet S1. Code is provided at https://github.com/hnishio/SIG5_field3.git.

