## Supplemental Figures and Tables for "Circadian and environmental signal integration in a natural population of *Arabidopsis*"

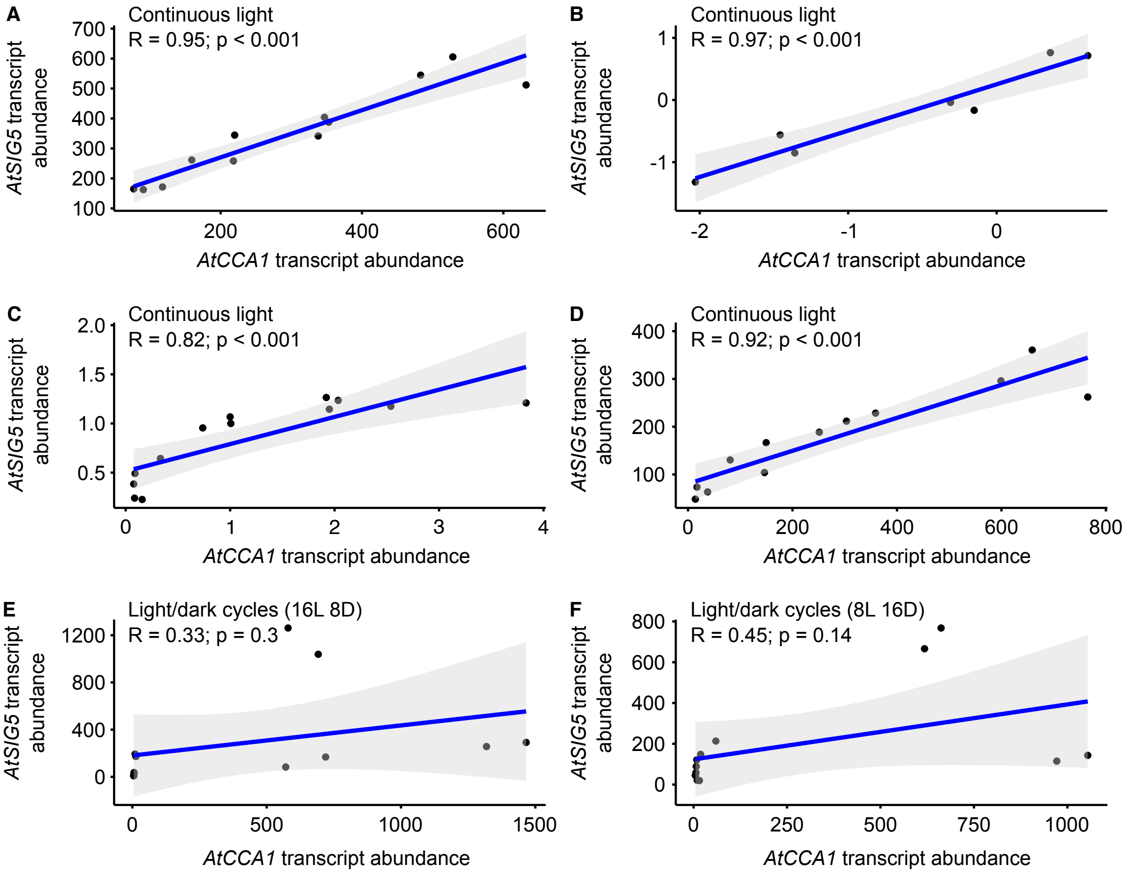


**Fig. S1.** Relationship between *AtCCA1* and *AtSIG5* transcript abundance in *A. thaliana* under controlled conditions. (A-D) Relationship between *AtCCA1* and *AtSIG5* transcript abundance under conditions of constant light, from the transcriptome studies of (A) [3] (B) [47], (C) [4], (D) [6]. (E, F) Relationship between *AtCCA1* and *AtSIG5* transcript abundance under light/dark cycles with (E) long and (F) short photoperiods, from the transcriptome study of [3, 48]. Blue lines indicate a regression line. Pearson's correlation coefficient (R) with p-values testing for the likelihood of a chance correlation are shown for each plot.

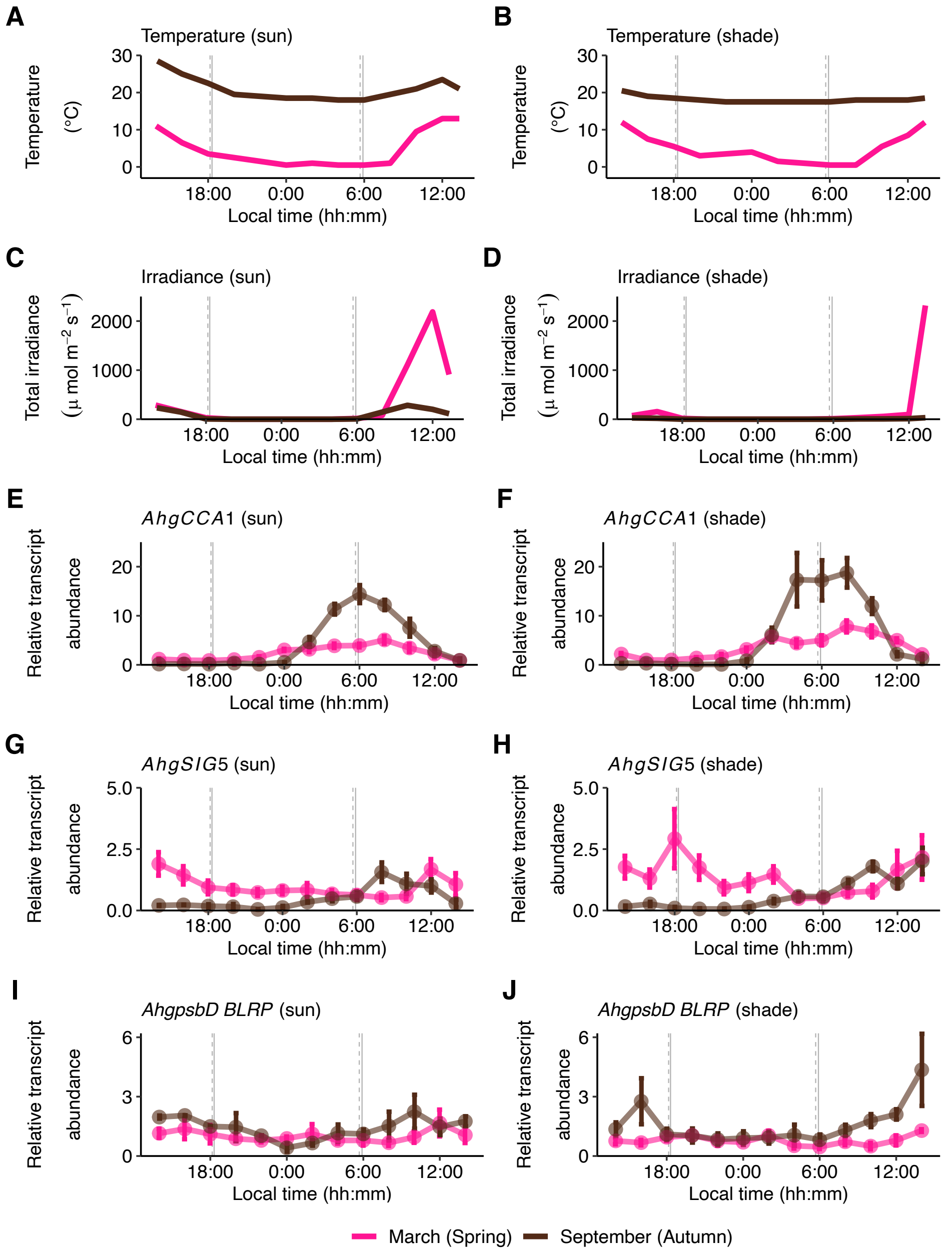


**Fig. S2.** Data underlying models produced in this study, here comparing signalling pathway dynamics between two seasons, under different light conditions in a natural population of *A. halleri*. (A-D) Diel fluctuations in (A, B) ambient temperature and (C, D) total irradiance detected (200-900 nm), at 2-hour intervals (thinned out from original data measured at 5-minute intervals, for the purpose of aligning intervals with the transcript data) during sampling period in March and September 2015. (E-J) Transcript abundance of (E, F) *AhgCCA1*, (G, H) *AhgSIG5* and (I, J) *AhgpsbD BLRP*. Vertical grey lines on time-series plots indicate the times of sunrise and sunset during March (solid line) and September (dashed). Data are mean ± s.e.m; $n = 6$ replicate plants.

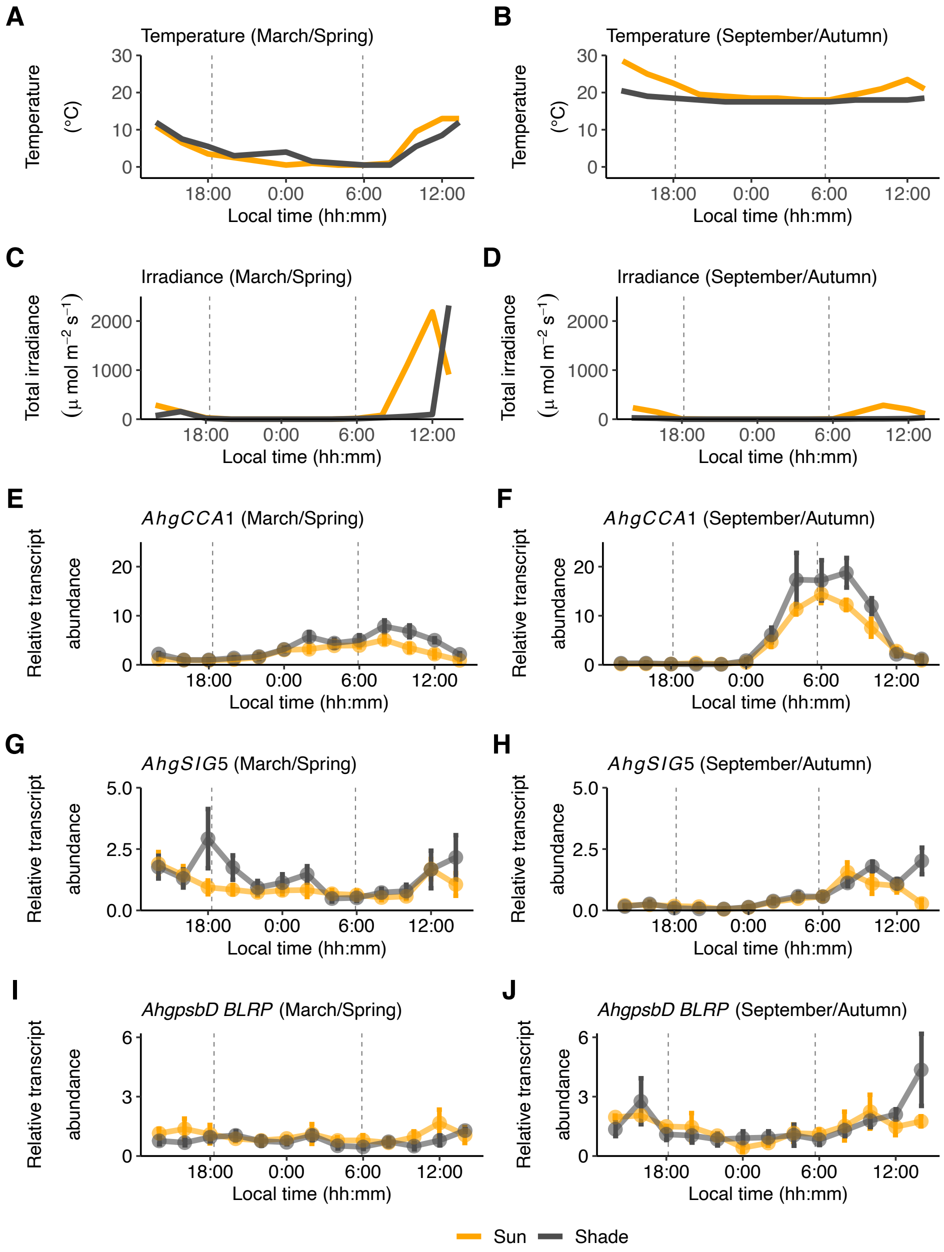


**Fig. S3.** Data underlying models produced in this study, here comparing signalling pathway dynamics between two different light conditions, during two sampling seasons in a natural population of *A. halleri*. (A-D) Diel fluctuations in (A, B) ambient temperature and (C, D) total irradiance detected (200-900 nm), at 2-hour intervals (thinned out from original data measured at 5-minute intervals, for the purpose of aligning intervals with the transcript data) during sampling period in March and September 2015. (E-J) Transcript abundance of (E, F) *AhgCCA1*, (G, H) *AhgSIG5* and (I, J) *AhgpsbD BLRP*. Vertical grey lines on time-series plots indicate the times of sunrise and sunset. Data are mean ± s.e.m; *n* = 6 replicate plants.

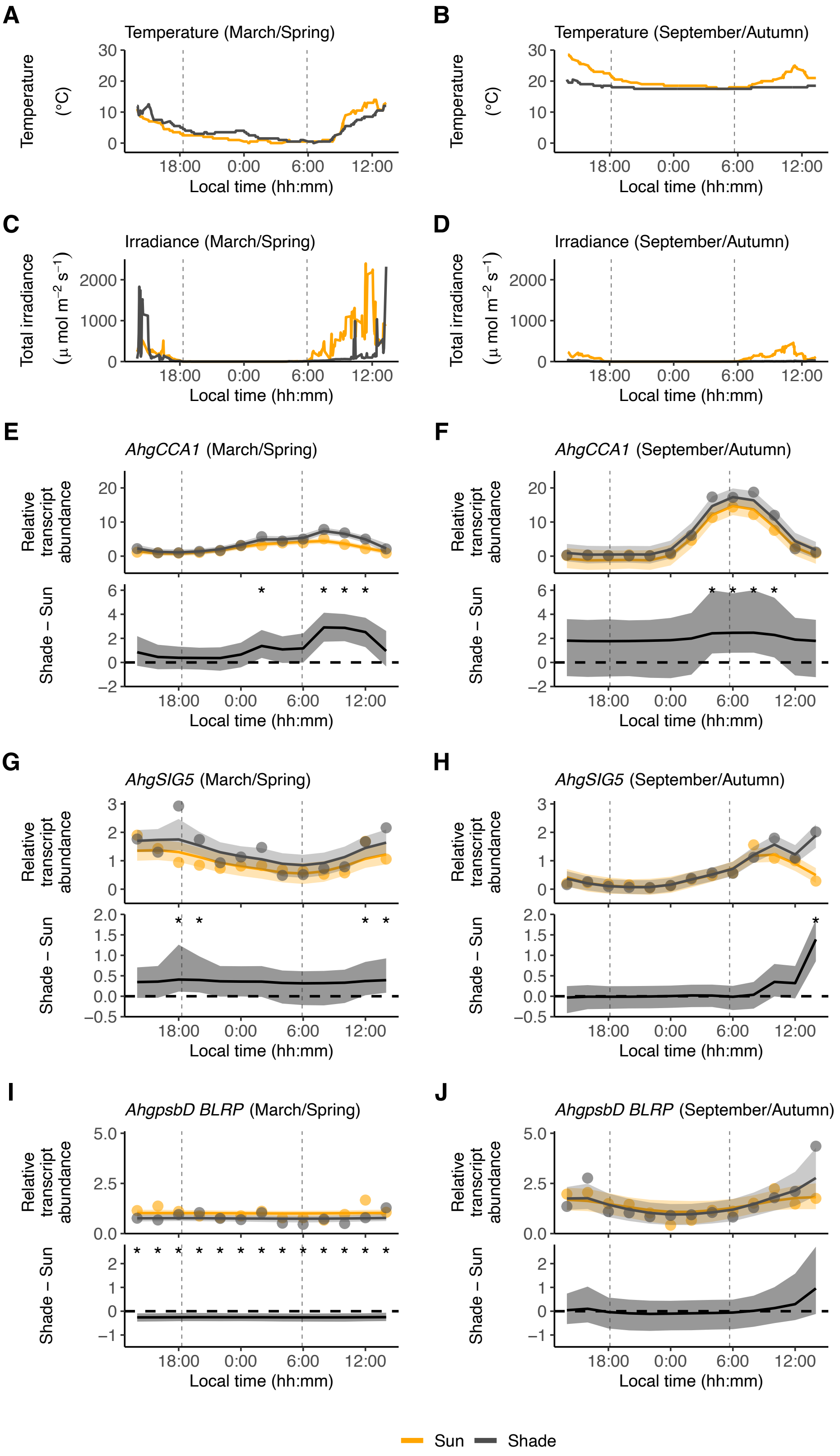


**Fig. S4.** Distinct diel dynamics of components of a circadian signalling pathway between sun and shade conditions in a natural population of *A. halleri*. (A-D) Diel fluctuations in (A, B) ambient temperature and (C, D) total irradiance detected (200-900 nm), measured at 5-

minute intervals during sampling period in March and September 2015. (E-J) Bayesian estimation of smooth trend model (STM) for transcript dynamics of (E, F) *AhgCCA1*, (G, H) *AhgSIG5* and (I, J) *AhgpsbD BLRP*. In each panel, the upper graphs show the predicted relative transcript abundance for sun (orange; $\mu_1$, equation 1 in Materials and methods) and shade (light grey; $\mu_2$, equation 3) conditions with the mean of observed values (dots), and the lower graphs represent the differences in transcript abundance between sun and shade conditions ($\delta$, equation 2). The solid line and the shaded region are the median and the 95% confidence interval of the posterior distribution. When the 95% confidence interval of the difference between sun and shade conditions does not contain zero, the difference is considered significant and is indicated by asterisks. Vertical grey lines on time-series plots indicate the times of sunrise and sunset. STM analysis used data from 6 replicate plants per condition.

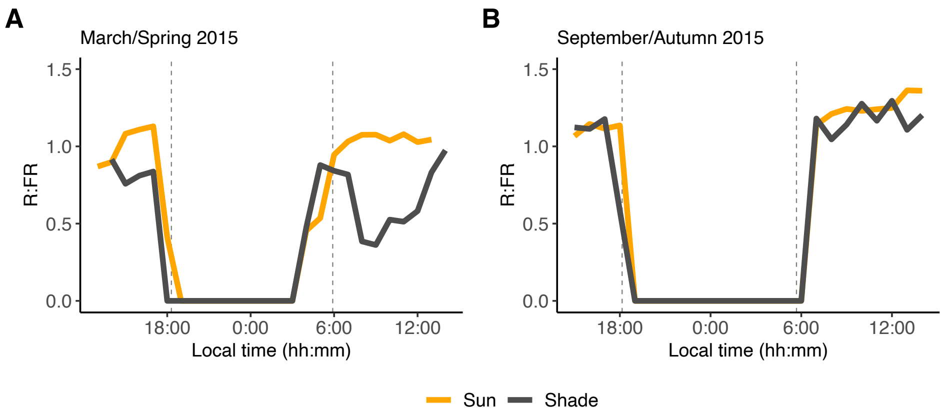


**Fig. S5.** The ratio of red to far-red light in a natural population of *A. halleri*, during March and September sampling seasons. (A, B) Comparison of the ratio of red to far-red light received by plants under the sun- and shade conditions during (A) March 2015 and (B) September 2015 sampling seasons. The R:FR varied during the photoperiod during both sampling seasons, and the effect of shade on R:FR was ameliorated by heavy cloud cover. Vertical grey lines on graphs indicate the times of sunrise and sunset.

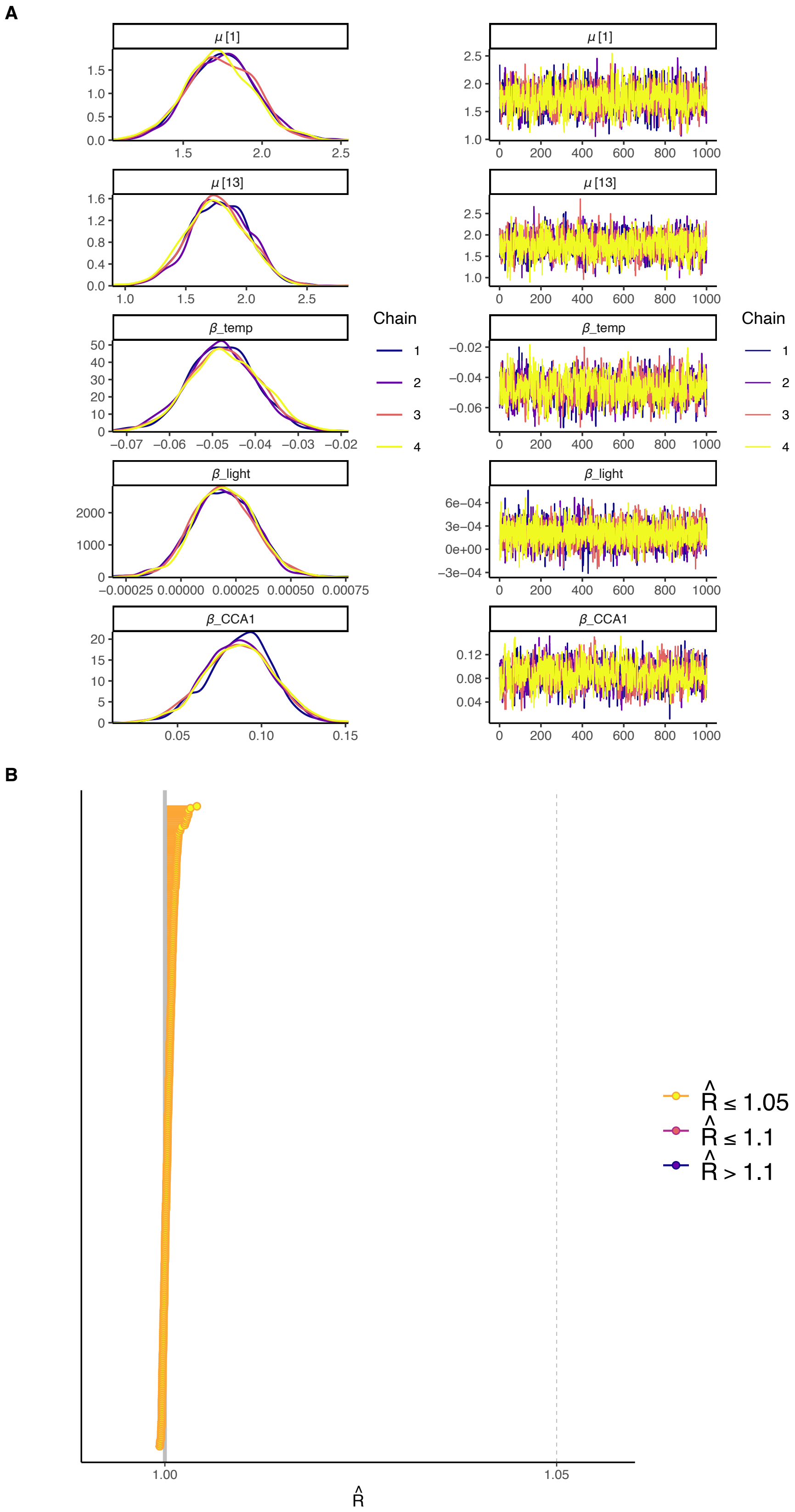


**Fig. S6.** Convergence of Markov Chain Monte Carlo (MCMC) sampling for local level model with exogenous variables (LLMX) for *AhgSIG5*. (A) Density plots (left) and trace plots (right)

of 1,000 MCMC samples/chain generated from posterior distributions of $\mu$ (auto regressive trend component) and $\beta$ (regression coefficients of temperature [temp], irradiance [light] and upstream gene [CCA1]), after 3,000 warm-up steps. $\mu[1]$ and $\mu[13]$ represent the trend component values at the first and last time points. (B) R-hat values (an indicator of how well chains are mixed, calculated by comparing within-chain variance and total variance) of all model parameters. R-hat ≤ 1.05 is recommended by the stan development team (https://mc-stan.org/rstan/reference/Rhat.html).

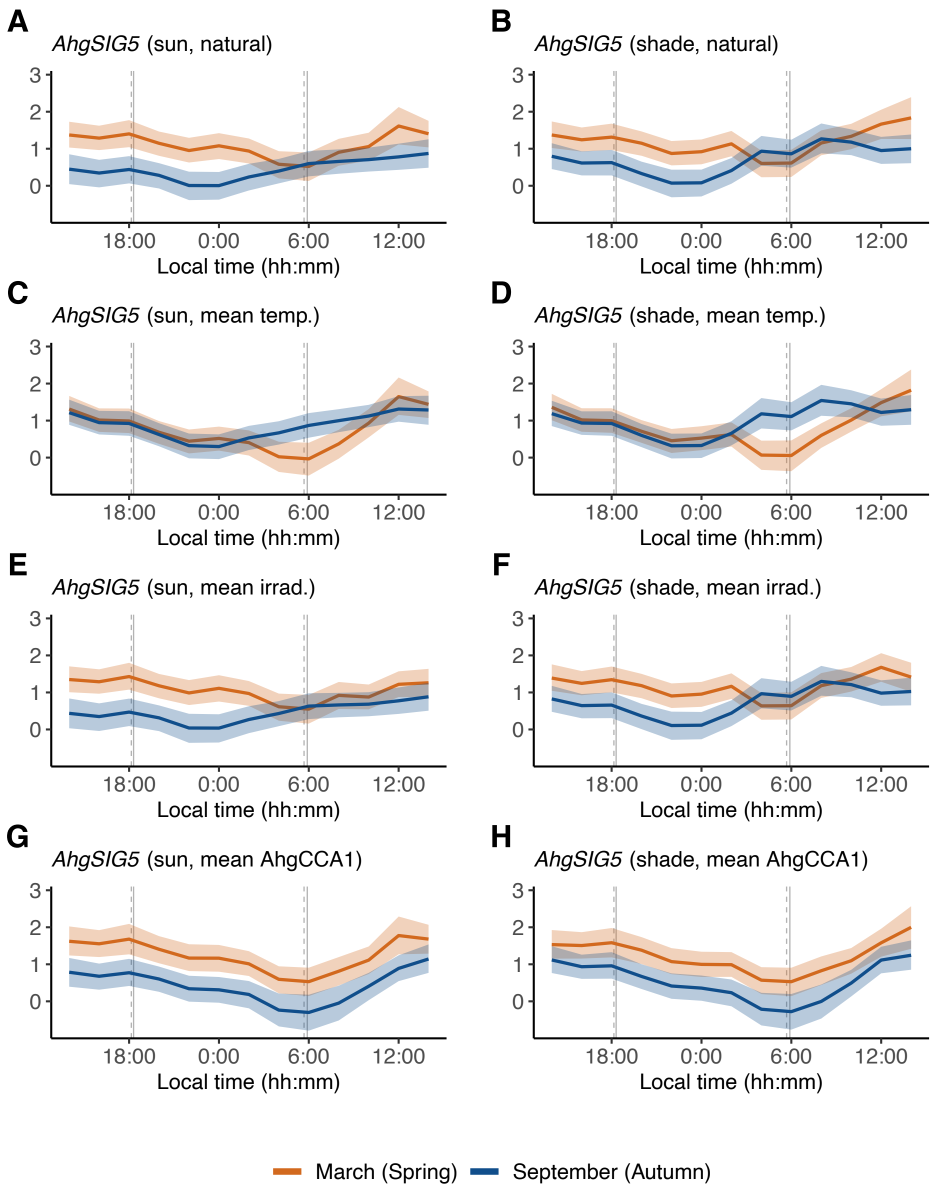


**Fig. S7.** Prediction of *AhgSIG5* transcript abundance using estimated parameter values in the local level model with exogenous variables (LLMX), when specific input variables are given as constant values. (A, B) LLMX prediction of *AhgSIG5* transcript abundance ($\alpha$, equations 10-13 in Materials and methods) where all variables are allowed to follow natural fluctuations (as in Fig. 3A, B). (C-H) LLMX prediction of *AhgSIG5* transcript abundance ($\alpha$) where (C, D) ambient temperature, (E, F) irradiance and (G, H) *AhgCCA1* transcript abundance were fixed at their mean value among all conditions. Shaded area represents 95% confidence interval. Vertical grey lines on time-series plots indicate the times of sunrise and sunset during March (solid line) and September (dashed).

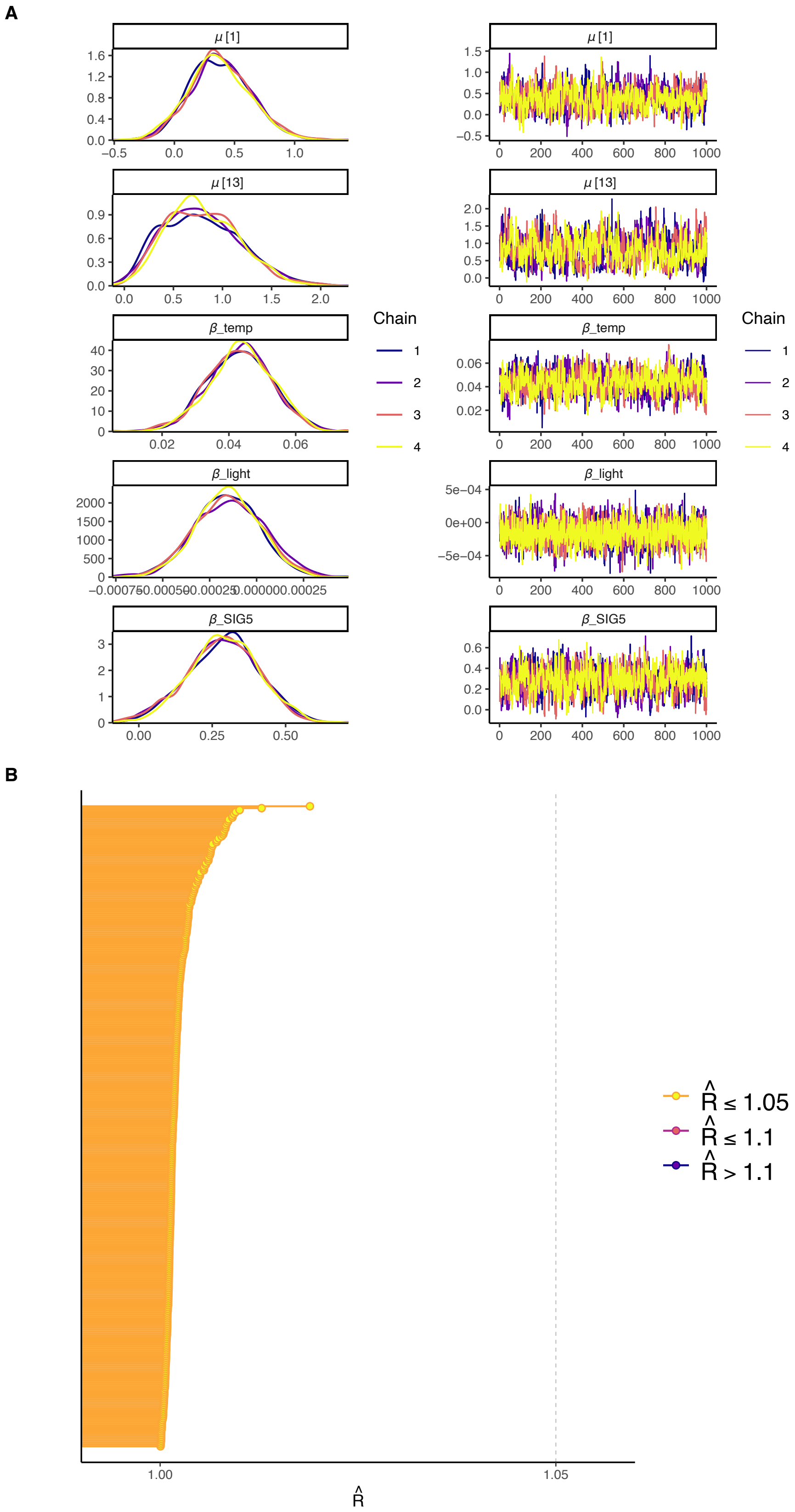


**Fig. S8.** Convergence of Markov Chain Monte Carlo (MCMC) sampling for local level model with exogenous variables (LLMX) for *AhgpsbD* BLRP. (A) Density plots (left) and trace plots (right) of 1,000 MCMC samples/chain generated from posterior distributions of $\mu$ (auto regressive trend component) and $\beta$ (regression coefficients of temperature [temp], irradiance [light] and upstream gene [SIG5]), after 3,000 warm-up steps. $\mu[1]$ and $\mu[13]$ represent the trend component values at the first and last time points. (B) R-hat values (an indicator of how well chains are mixed, calculated by comparing within-chain variance and total variance) of all model parameters. R-hat ≤ 1.05 is recommended by the stan development team (https://mc-stan.org/rstan/reference/Rhat.html).

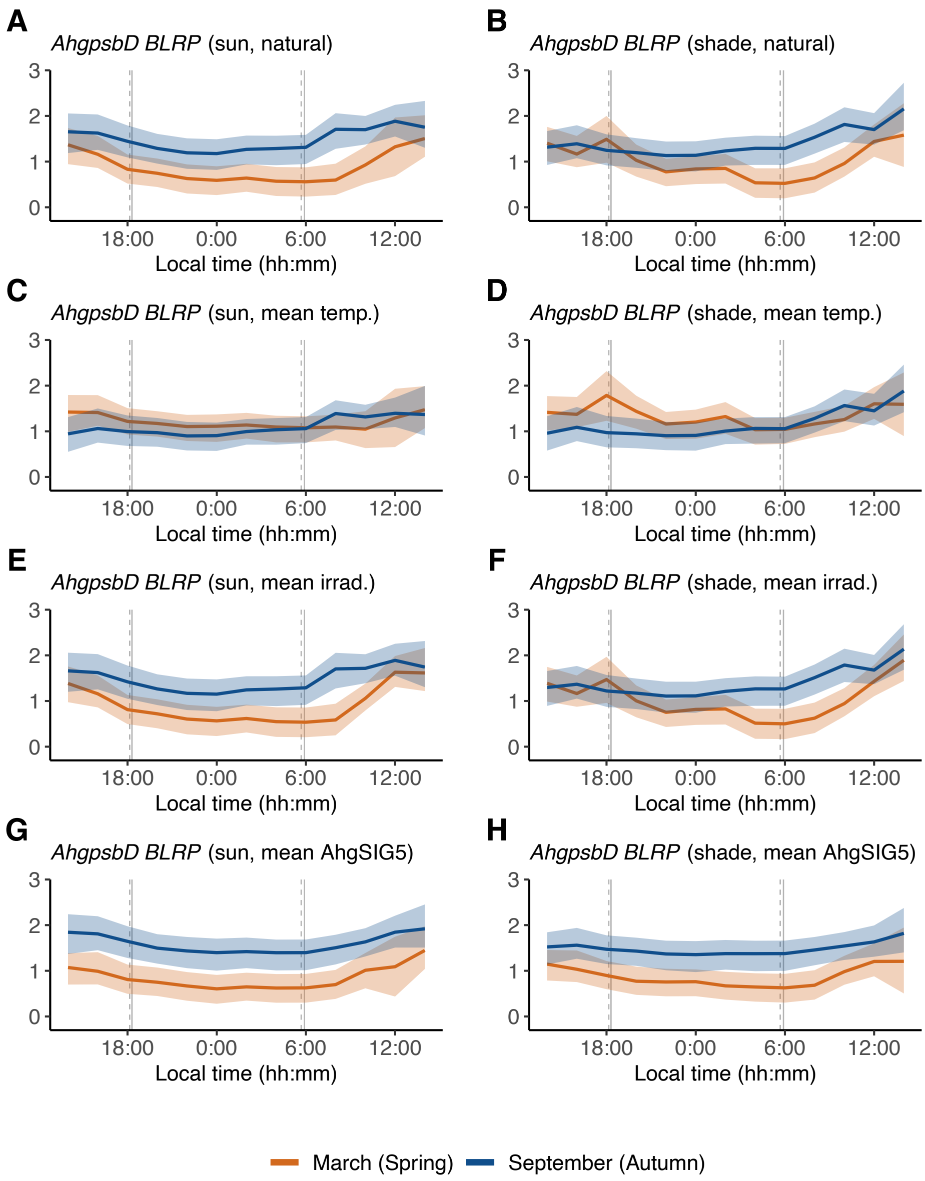


**Fig. S9.** Prediction of *AhgpsbD* BLRP transcript abundance using estimated parameter values in the local level model with exogenous variables (LLMX), when specific input variables are given as constant values. (A, B) LLMX prediction of *AhgpsbD* BLRP transcript abundance ($\alpha$, equations 10-13 in Materials and methods) where all variables are allowed to follow natural fluctuations (as in Fig. 3E, F). (C-H) LLMX prediction of *AhgpsbD* BLRP transcript abundance ($\alpha$) where (C, D) ambient temperature, (E, F) irradiance and (G, H) *AhgSIG5* transcript abundance were fixed at their mean value among all conditions. Shaded area represents 95% confidence interval. Vertical grey lines on time-series plots indicate the times of sunrise and sunset during March (solid line) and September (dashed).

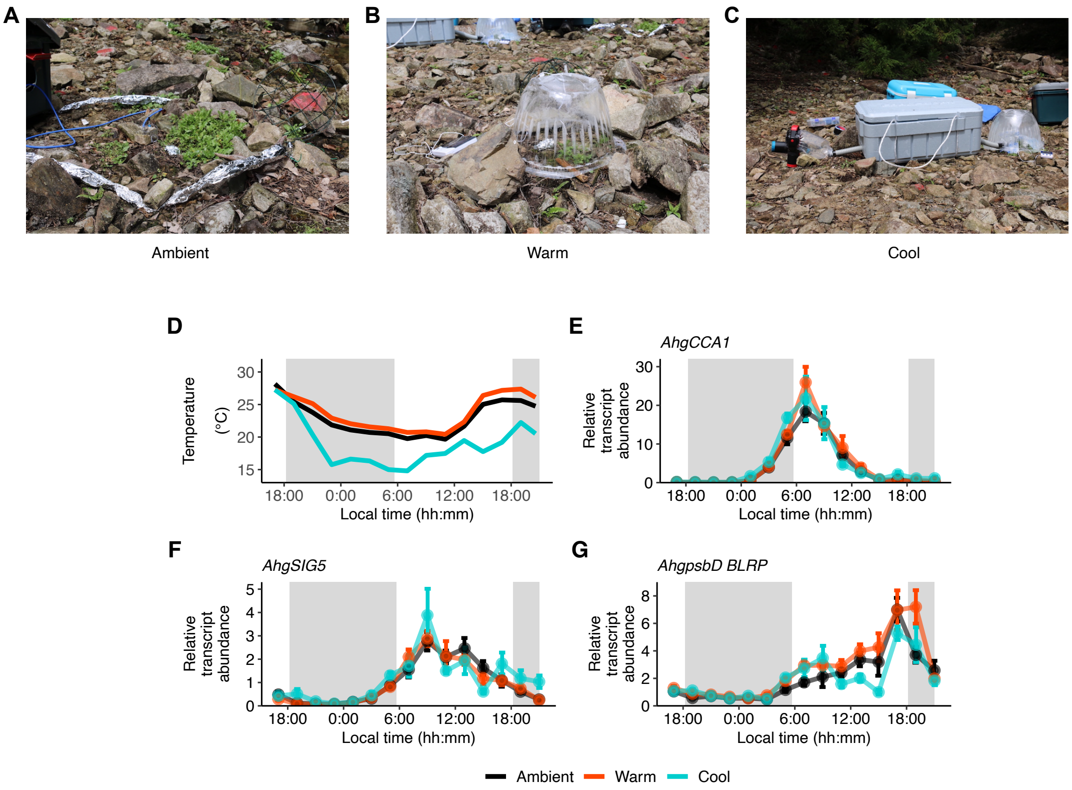


**Fig. S10.** Moderate temperature manipulations applied to patches of *A. halleri* plants, in the field, using custom-designed equipment. (A) Representative appearance of plant patches under naturally fluctuating conditions. (B) Plants covered with a plastic dome to raise the temperature. (C) Plants covered with a plastic dome undergoing temperature reduction with a custom chilling device. In this device, cool air is introduced to enclosed plant patches after being driven slowly through a heat exchanger, positioned within an expanded polystyrene box filled with ice. (D) Temperature changes in each condition at 2-hour intervals (thinned out from original data measured at 5-minute intervals, for the purpose of aligning intervals with the transcript data) during sampling period in September 2016. (E-G) Fluctuations in *AhgCCA1*, (F) *AhgSIG5* and (G) *AhgpsbD* BLRP transcript abundance under ambient conditions and following temperature manipulation of plant patches. Grey shaded boxes on graphs indicate the period between sunset and sunrise. Data are mean ± s.e.m; $n$ = 6 replicate plants.

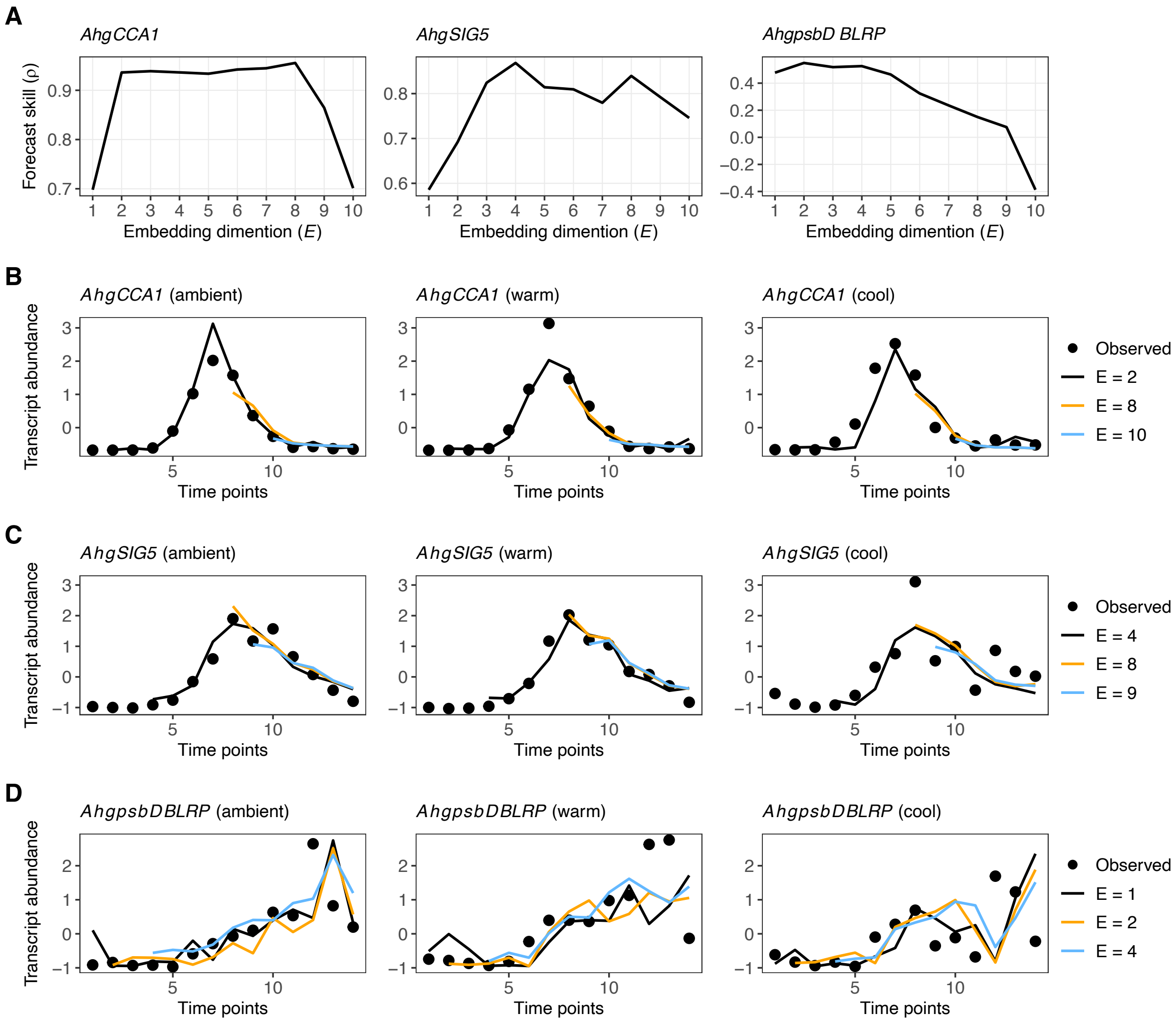


**Fig. S11.** Determination of optimal embedding dimensions of components of a circadian signalling pathway for temperature manipulation experiments in September 2016. (A) Evaluation of optimal embedding dimension *E* for each pathway component, with the forecast skill measured using ρ. A greater forecast skill was used to select the most appropriate embedding dimension. (B-D) Prediction of the fluctuations of *AhgCCA1*, *AhgSIG5* and *AhgpsbD* BLRP for several *E* values. Dots and lines are the observed (mean of replicates) and predicted values, respectively. The optimal *E* values determined were 2 for *AhgCCA1*, 4 for *AhgSIG5*, and 2 for *AhgpsbD* BLRP (see Materials and methods for more detail).

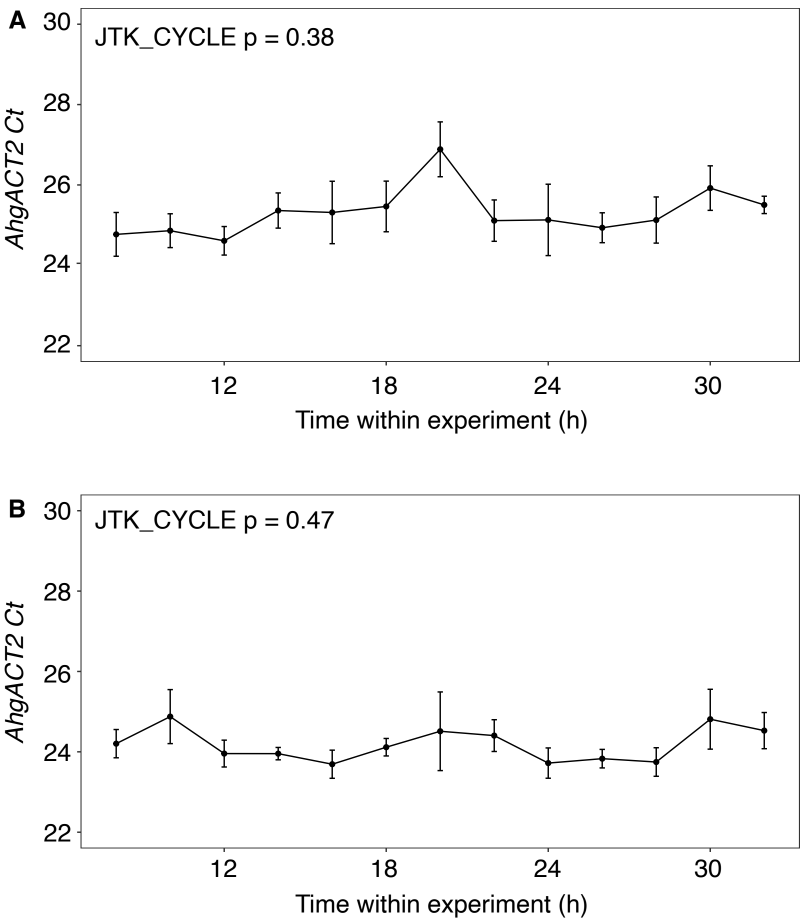


**Fig. S12.** *AhgACT2* reference transcript was not rhythmic over diel timecourses in our experiments. Ct for *AhgACT2* from RT-qPCR analysis of (A) sun and (B) shade datasets collected during September 2015. Data are mean Ct ± s.e.m; $n = 6$. p-values provided for statistical test for significant rhythmicity, using JTK_CYCLE algorithm [80].

| Primer | Sequence (5’ to 3’) |
|---|---|
| *AhgACT2* (forward) | TCAGATGCCCAGAAGTGTTGTTCC |
| *AhgACT2* (reverse) | CCGTACAGATCCTTCCTGATATCC |
| *AhgpsbD* BLRP (forward) | GGAAATCCGTCGATATCTCT |
| *AhgpsbD* BLRP (reverse) | CTCTCTTTCTCTAGGCAGGAAC |
| *AhgSIG5* (forward) | GTGTTGGAGCTAATAACAGCAGACA |
| *AhgSIG5* (reverse) | TGTCGAATAACCAGACTCTCTTTCG |
| *AhgCCA1* (forward) | GCACTTTCCGCGAGTTCTTG |
| *AhgCCA1* (reverse) | TGACTCCTTTCTTATCCTGTTATTCTG |

**Table S1.** Primers used for RT-qPCR analysis of transcript abundance in *Arabidopsis halleri* subsp. *gemmifera*.

| Parameter | 2.5% | 50.0% | 97.5% |
|---|---|---|---|
| μ[1] | 1.32 | 1.74 | 2.16 |
| μ[2] | 1.13 | 1.48 | 1.85 |
| μ[3] | 1.11 | 1.49 | 1.87 |
| μ[4] | 0.83 | 1.17 | 1.50 |
| μ[5] | 0.56 | 0.89 | 1.23 |
| μ[6] | 0.49 | 0.84 | 1.16 |
| μ[7] | 0.35 | 0.71 | 1.08 |
| μ[8] | -0.20 | 0.27 | 0.72 |
| μ[9] | -0.30 | 0.20 | 0.68 |
| μ[10] | 0.02 | 0.50 | 0.98 |
| μ[11] | 0.58 | 0.99 | 1.41 |
| μ[12] | 1.19 | 1.61 | 2.06 |
| μ[13] | 1.29 | 1.76 | 2.24 |
| β_temp | -0.06 | -0.05 | -0.03 |
| β_light | -1.00E-04 | 2.00E-04 | 5.00E-04 |
| β_CCA1 | 0.05 | 0.09 | 0.13 |
| σ_μ | 0.23 | 0.41 | 0.72 |
| σ_Y | 0.77 | 0.84 | 0.91 |

**Table S2.** Estimated parameter values in the local level model with exogenous variables (LLMX) for *AhgSIG5*. The 2.5 %, 50.0 % and 97.5 % points of 4,000 MCMC samples obtained from posterior distributions are shown.

| Parameter | 2.5% | 50.0% | 97.5% |
|---|---|---|---|
| μ[1] | -0.09 | 0.37 | 0.90 |
| μ[2] | 0.04 | 0.48 | 1.07 |
| μ[3] | -0.01 | 0.41 | 0.89 |
| μ[4] | 0.00 | 0.39 | 0.83 |
| μ[5] | -0.03 | 0.35 | 0.76 |
| μ[6] | -0.04 | 0.33 | 0.73 |
| μ[7] | 0.00 | 0.35 | 0.76 |
| μ[8] | -0.03 | 0.35 | 0.74 |
| μ[9] | -0.02 | 0.35 | 0.75 |
| μ[10] | 0.02 | 0.41 | 0.85 |
| μ[11] | 0.08 | 0.50 | 1.03 |
| μ[12] | 0.10 | 0.60 | 1.22 |
| μ[13] | 0.15 | 0.76 | 1.58 |
| β_temp | 0.02 | 0.04 | 0.06 |
| β_light | -5.00E-04 | -2.00E-04 | 2.00E-04 |
| β_SIG5 | 0.04 | 0.29 | 0.52 |
| σ_μ | 0.02 | 0.16 | 0.45 |
| σ_Y | 1.00 | 1.08 | 1.18 |

**Table S3.** Estimated parameter values in the local level model with exogenous variables (LLMX) for *AhgpsbD* BLRP. The 2.5 %, 50.0 % and 97.5 % points of 4,000 MCMC samples obtained from posterior distributions are shown.
